# Metabolic Flexibility of Microglia: Energy Substrate Utilization and Impact on Neuronal Metabolism

**DOI:** 10.1101/2025.09.18.676073

**Authors:** Emil W. Westi, Rebecca Birch Carlsen, Jens Velde Andersen, Zeliha Kilic, Dana Alnajar, Nadia K Holmgaard, Belén García Sintes, Agustin Cota-Coronado, Sonia Sanz Muñoz, Céline Galvagnion, Blanca I. Aldana

**Author notes:** Correspondence:* Blanca I. Aldana -, Postal address: Department of Veterinary and Animal Sciences, PAP Grønnegårdsvej 7, Building 1-04, Room T01, 1870 Frederiksberg C, Denmark.

## Abstract

Microglia, the main resident immune cells of the brain, play critical roles in maintaining neuronal function and homeostasis. Microglia metabolic flexibility enables rapid adaptation to environmental changes, yet the full extent of their metabolic capabilities and influence on neuronal metabolism remains unclear. While microglia predominantly rely on glucose oxidative metabolism under homeostatic conditions, they shift toward glycolysis upon proinflammatory activation. In this study, we investigated microglial metabolism and its impact on neuronal metabolic homeostasis using isotope tracing with stable carbon ^13^C-enriched substrates and gas chromatography-mass spectrometry (GC-MS) analysis. Primary microglia were incubated with ^13^C-labeled glucose, glutamine, or GABA in the presence or absence of lipopolysaccharide (LPS) to assess metabolic adaptations upon an inflammatory challenge. Additionally, neurons co-cultured with quiescent or activated microglia (either with LPS or amyloid-β) were incubated with ^13^C-enriched glucose to examine microglia-neuron metabolic interactions. Our findings confirm that microglia readily metabolize glucose and glutamine, with LPS stimulation slightly changing the glycolytic activity, as indicated by subtle changes in extracellular lactate. Importantly, we demonstrate for the first time that microglia take up and metabolize the inhibitory neurotransmitter GABA, suggesting a novel metabolic function. Furthermore, microglial presence directly influences neuronal metabolism and neurotransmitter homeostasis, highlighting a previously unrecognized aspect of neuron-microglia metabolic crosstalk. Collectively, these findings provide new insights into microglial metabolism and its role in neuronal function, with implications for neuroinflammatory and neurodegenerative diseases in which microglial metabolism is dysregulated.

## Introduction

Microglia, the innate immune cell of the central nervous system (CNS), are essential for neuronal and synaptic function (Nayak, Roth, & McGavern, 2014). Microglia serve as the first line of defense against foreign pathogens and injuries (Hanisch & Kettenmann, 2007; Kettenmann et al., 2011). During quiescent states, microglia are highly active with ramified processes constantly surveying the brain parenchyma by dynamically extending and retracting (Nimmerjahn, Kirchhoff, & Helmchen, 2005; Wake et al., 2009). In recent years, it has become evident that microglia constitute a highly heterogeneous cell population, comprising multiple subtypes even under homeostatic conditions (Stratoulias et al., 2019). This allows microglia to quickly adapt to alterations in the immediate environment, changing morphology and cellular functions accordingly (Butovsky & Weiner, 2018). Similar to peripheral immune cells, microglia are required to maintain cellular functions even under challenging energetic conditions. To achieve this, peripheral immune cells can metabolize different auxiliary energy substrates in addition to glucose (Buck, O’Sullivan, & Pearce, 2015; Caputa, Castoldi, & Pearce, 2019; Van den Bossche, O’Neill, & Menon, 2017). This metabolic flexibility is particularly important as it enables immune cells to carry out functions in disrupted environments with varying access to glucose. Exposure to the bacterial endotoxin lipopolysaccharide (LPS), mimicking an inflammatory response, leads to a shift from mitochondrial oxidative phosphorylation towards glycolytic activity in macrophages, being a necessary step in the proinflammatory activation (Viola et al., 2019). Surveilling microglia express similar metabolic characteristics as macrophages, relying on oxidative phosphorylation of glucose, the primary fuel of the brain, but shifting towards glycolysis upon exposure to proinflammatory stimuli (Gimeno-Bayón et al., 2014; Nair et al., 2019; Voloboueva et al., 2013; York et al., 2021). However, when glucose is scarce, microglia are also able to rely on oxidation of glutamine and fatty acids (Bernier et al., 2020). Likewise, it has been observed, *in vitro*, that microglia will increase GLT-1 expression following LPS-stimulation (Persson et al., 2005). The metabolism of these alternative substrates may even be crucial for microglia function under surveilling conditions (Bernier et al., 2020). Although microglia share similar roles with peripheral macrophages, the energetic demands within the distinct metabolic environment of the brain are far from elucidated. So far, most studies have assessed microglia metabolism using indirect measurements (Bernier et al., 2020; York et al., 2021). Additionally, despite extensive research on microglia, little is known about the interplay between microglia and neuronal metabolism. The overall aim of this study was therefore to assess functional microglial metabolism and to explore potential effects of microglial function on neuronal metabolism. To achieve this, we incubated primary cultures of microglia with energy substrates enriched with the stable carbon isotope ^13^C in the presence or absence of LPS. Primary cultures of neurons co-cultured with quiescent or stimulated microglia were additionally incubated with ^13^C-enriched substrates. The intracellular isotopic enrichment of metabolites was assessed by gas chromatography-mass spectrometry (GC–MS) analysis. We report that primary cultures of microglia readily take up and metabolize glucose and glutamine. In addition, lactate amounts released to the media were only slightly increased, when microglia are stimulated with LPS. We demonstrate, for the first time, that microglia are able to take up and metabolize the inhibitory neurotransmitter GABA and that the presence of microglia directly affects neuronal metabolism and neurotransmitter homeostasis. Overall, our results provide new insights into functional metabolism of microglia and their metabolic communication with neurons, which will be central to understanding the functional role of microglia in both health and disease.

## Methods

### Animals and ethical approval

Mouse breeding pairs (NMRI; RRID:IMSR_ENV:HSD-275) were purchased from Envigo (Horst, Netherlands) and housed at the Department of Drug Design and Pharmacology, University of Copenhagen, in a specific pathogen-free animal facility with humidity and temperature-control. The mice were kept under a 12-hour light/dark cycle with free access to water and chow. Both female and male pups were used at 7-days of age to prepare primary cultures of microglia. Experiments were approved by the Danish National Ethics Committee and performed according to the European Convention (ETS 123 of 1986). The experiments using animals have been reported in compliance with the ARRIVE guidelines.

### Materials

The stable ^13^C enriched compounds [U-^13^C]glucose (CLM-1396-5, 99%), [U-^13^C]glutamine (CLM-1822-H-PK, 99%) and [U-^13^C]GABA (CLM-8666, 98%) were all purchased from Cambridge Isotope Laboratories (Tewksbury, USA). Dulbecco’s modified Eagle’s medium (DMEM) powder, poly-D-lysine, N-tert-butyldimethylsilyl-N-methyltrifluoroacetamide (MTBSTFA), N,N-dimethylformamide (DMF) and DAPI (ready-made solution), lipopolysaccharide from Escherichia coli O111:B4 (LPS, Sigma L2630), were purchased from Sigma-Aldrich (St. Louis, MO, USA). 25 cm^2^ culturing flasks and 6-well plastic culture dishes were purchased from NUNC A/S (Roskilde, Denmark). Rabbit anti-IBA1 polyclonal antibody (019-19741,RRID:AB_839504) was purchased from Fujifilm (Osaka, Japan). Chicken anti-GFAP polyclonal antibody (PA1-10004, RRID:AB_1074620) and goat anti-chicken Alexa Fluor 594 (A-11042; RRID:AB_2534099) were obtained from Thermo Fisher Scientific (Waltham, MA, USA). Donkey anti-rabbit Alexa Fluor 488 (711-545-152; RRID:AB_2313584) was purchased from Jackson ImmunoResearch Laboratories (Cambridge House, Britain). Culture 6-well Plate (GR-657160) and ThinCerts 6m/PET membrane Pore O = 0.4 µm (GR-657641) from Greiner Bio-One (Austria). Other chemicals used were of the purest grade available from regular commercial sources.

### Preparation of primary cultures of microglia

The handling and preparations of the primary microglia cell cultures was done under aseptic condition in a laminar air flow cabinet. All solutions were sterile filtered using a PALL AcrocapTM with 0.2 μm Supor membrane (Pall corporation, NY, USA) or sterile syringe filters with 0.2 μm polyethersulfone membranes (VWR International, PA, USA) prior to use. Primary cultures of glial cells were prepared and cultured according to Walls et al. (Anne B. Walls et al., 2014). Briefly, 7-day old NMRI pups were decapitated, and cortices were dissected, pooled and placed on nylon mesh (80 µm pore size) in a petri dish containing 10 mL DMEM-based culture medium (reconstituted DMEM powder (Sigma, D5030)) supplemented with 2.5 mM L-glutamine, 6 mM D-glucose, 26.2 mM NaHCO_3_, 100 IU/mL penicillin and 0.04 mM phenol red. While DMEM differs from the ionic composition and nutrient levels found in native brain CSF or ECF, its use here reflects established practices in glial and neuronal primary culture (Walls et al., 2014; Lian et al., 2016). DMEM supports the metabolic activity and viability necessary for isotope tracing experiments. Upon which, mechanical dissociation was executed by squeezing the cortices through the mesh using a 20 mL syringe. The suspension was then triturated 3 times using a syringe fitted with a steel canula (13G), prior to seeding the cells in an appropriate density in 25 cm^2^ flasks. The primary cultures were kept at 37°C and 95% atmospheric air and 5% CO_2_ for two weeks and media was exchanged twice a week. Following two weeks of culturing the microglia were separated from the astrocyte monolayer as described in Lian et al. (2016) (Lian, Roy, & Zheng, 2016). The flasks were gently tapped under aseptic conditions and the supernatant containing the microglia was gently removed as to not disturb the astrocyte monolayer. The supernatant was centrifuged at 0.3 *g* and the supernatant was aspirated and the pellet redissolved in conditioned media and seeded in 6-well plates. Prior to seeding, 6-well plates were coated with poly-D-lysine and washed with Tyrodes Buffer (136.9 mM NaCl, 0.57 mM NaHPO_4_ and 0.36 nM NaH_2_PO_4_, pH 7.4). Following two hours after seeding, the microglia were visually inspected under a microscope to assure proper adherence to the bottom and the media was exchanged. In certain data sets, primary cultures of microglia were exposed to LPS (100 ng/mL) for 24h to induce activation.

### Preparation of primary culture of cortical neurons

Cerebral cortical neurons were isolated from 14-day NMRI mouse fetuses and cultured as previously described (Anne B. Walls et al., 2014). The pregnant mice (one dam per cell preparation indicated as N in the corresponding figure) were killed by cervical dislocation, fetuses (8-12 fetuses per dam) were dissected into a Petri dish and each fetus is separated from the amnion and decapitated to collect and pool the cortices for each cell preparation. Cells were seeded in poly-D-lysine coated 6-well plates (washed with Tyrode’s buffer) in a modified DMEM containing 10% fetal bovine serum (FBS). This culture medium included reconstituted DMEM powder, 19 mM KCl, 12 mM D-glucose, 26.2 mM NaHCO₃, 0.2 mM L-glutamine, 0.1 IU/L insulin, 7.3 µM p-aminobenzoic acid, and 100 IU/mL penicillin. After 36–48 hours, 20 µM cytosine arabinoside was added to prevent glial proliferation. Neuronal cultures were maintained for 7 days, with D-glucose added on days two and five. It has been reported that primary cultures from E14 mouse cerebral cortex are typically considered to have approximately 20% GABAergic neurons due to rapid developmental progression and in vitro context (Sahara et al., 2012).

### Non-contact (NC) co-culture model

The co-culture system was adapted from Roque et al. 2017 (Roqué & Costa, 2017) and consisted of permeable culture inserts containing the microglia cells and cell culture 6-well plates seeded with the primary cultures of neurons. Primary cultures of glia were plated and incubated for two weeks in 25 cm^2^ flasks. After this time, the microglia cultures were isolated and plated in the permeable culture inserts (pore size 0.4 μm, ThinCert 657641) for 24h. These primary cultures of cortical neurons were plated in the 6-well plates and kept for one week at 37°C. The co-cultures were assembled and kept for two hours under the different experimental conditions. For some experimental sets, to induce activation, microglia were stimulated with either LPS (100 ng/mL) or 500 nM amyloid-beta (Aβ)_1-42_ for 24 hours prior to co-culturing with neurons. This approach was also designed to prevent direct exposure of neurons to LPS or Aβ. Cultures of primary cultures of neurons (monocultures) were used as control.

### Cell Viability Assay

To assess the effect of microglia on neuronal survival, the tetrazolium salt XTT Assay (Sigma, X4626) was employed. Briefly, after co-culturing with microglia, microglia inserts were removed and the media in the neuronal wells were aspirated. Assay media containing DMEM (without phenol red), 1 mg/mL XTT, 5 μM N-methylphenazonium methyl sulfate (PMS), and 2.5 mM D-glucose was added to each well and incubated for three hours in the dark at 37°C. Media samples from each well were transferred to a 96-well plate, and absorbance was measured at 450 nm and 650 nm using a spectrophotometer. Cell viability was assessed by subtracting the absorbance at 650 nm from 450 nm to correct for background absorbance. The absorbance of the XTT media was subtracted from each sample to isolate the absorbance specific to viable cells. Data were normalized to the monoculture designated as 100%.

### Incubation of primary cultures of microglia and neurons using stable ^13^C substrates

On the day of the experiment primary cultures of microglia were examined under microscope to assess confluency and morphology. After which, the culturing medium was removed by aspiration from the cell cultures, and the cells were washed once with 1 mL 37°C phosphate-buffered saline (PBS; 137 mM NaCl, 2.7 mM KCl, 7.3 mM Na_2_HPO_4_, 1.5 mM KH_2_PO_4_, 0.9 mM CaCl_2_ and 0.5 mM MgCl_2_, pH 7.4). The microglia cultures were incubated for 90 minutes at 95% atmospheric air/5% CO_2_ 37°C in 2 mL incubation media (serum-free DMEM, 26.2 mM NaHCO_3_) and different ^13^C-labeled substrates. The primary cultures of microglia were incubated with either 2.5 mM [U-^13^C]glucose, 0.5 mM [U- ^13^C]glutamine or 200 µM [U-^13^C]GABA in the presence or absence 100 ng/mL LPS. All conditions, except [U-^13^C]glucose, were supplemented with 2.5 mM unlabeled D-glucose. After 90 minutes, 1.5 mL of incubation media were stored for later analysis and the rest of the media was removed by aspiration. The cells were washed once in 1 mL ice cold PBS and extracted using ethanol (70% v/v), while on ice. To achieve sufficient amounts of biological material to detect ^13^C enrichment, the contents of two wells were pooled, yielding three samples per 6-well plate. After extraction the samples were centrifuged for 20 minutes (20,000 *g*, 4°C), thereby separating the extract (supernatant) from the pellet. The supernatant was lyophilized, reconstituted in H_2_O, and stored for later analyses. The pellets were reconstituted in 150 µL 1M KOH and for the amount of protein was determined.

In the non-contact (NC) co-culture system, the extracellular media in the neuronal cultures were aspirated and washed with 37°C PBS before adding incubation media containing 2.5 mM D-glucose (37°C). Hereafter, the extracellular media in the microglia-containing inserts were aspirated and washed with 37°C PBS. Each insert was transferred to their respective “neuronal” well, and incubation media containing 2.5 mM D-glucose (37°C) was added to each insert. The co-cultures and the monoculture were incubated for one hour. The extracellular media in the neuronal wells were aspirated, and incubation medium (37°C) containing 2.5 mM [U-^13^C]glucose was added to the wells. The microglia inserts were also aspirated, and DMEM medium (without labeled substrates or glucose) was added to each insert. The co-culture and the monoculture were incubated for one hour (37°C). After this time, inserts were discarded and the extracellular medium from the neurons were collected and stored at -20°C for later analyses. Neuronal cells were washed once in 1 mL ice cold PBS and extracted directly using ethanol (70% v/v), while on ice. After extraction the samples were centrifuged for 20 minutes (20,000 *g*, 4°C), thereby separating the extract (supernatant) from the pellet. The supernatant was lyophilized, reconstituted in H_2_O, and stored for later analyses. The pellets were reconstituted in 150 µL 1M KOH and used for protein determination.

### Dynamic metabolic mapping by ^13^C isotope tracing using gas chromatography-mass spectrometry (GC-MS)

Determination of ^13^C enrichment in tricarboxylic acid (TCA) cycle intermediates and connected amino acids in aqueous extracts was done as previously described in Westi et al. (Westi et al., 2022). The lyophilized samples were reconstituted in 150 µL of H_2_O, acidified with 1 M HCl and extracted twice with ethanol (96% v/v). Metabolites were derivatized utilizing MTBSTFA and the samples were analyzed by GC (RRID:SCR_019445 Agilent Technologies, 7820A, J&W GC column HP-5 MS) coupled to MS (RRID:SCR_019420, Agilent Technologies, 5977E). Isotopic enrichment was corrected for natural abundance of ^13^C as described in Walls et al. (Anne B. Walls et al., 2014). Data from incubations with [U-^13^C]glucose is displayed as M+X, where M is the molecular ion and X denotes the number of ^13^C atoms in the molecule. In addition, the molecular carbon labeling (MCL), the weighted average percentage of all the isotopologues of a metabolite, provides a measurement of the overall ^13^C accumulation (Andersen, Christensen, et al., 2021). Data from incubations with [U-^13^C]glutamine and [U-^13^C]GABA is presented as M+X, illustrating both direct and first turn metabolism of the substrates (Andersen et al., 2017; Andersen et al., 2024). Data from the NC set-up are presented as either M+X or MCL.

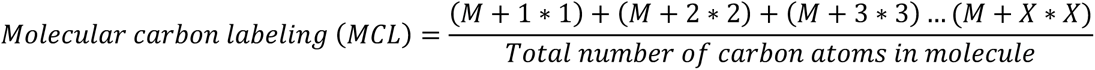

Due to the distinct entry points of the different ^13^C-labeled substrates into the TCA cycle, the resulting variability in metabolite labeling patterns and detectability, combined with the focus on biologically relevant signals and cell-type–specific metabolic pathways, only those metabolites with consistent ^13^C enrichment above background are selectively reported to ensure relevance in each experimental condition.

### Quantification of amino acids by High Performance Liquid Chromatography (HPLC)

The reconstituted aqueous cell extracts were analyzed by reverse-phase HPLC (RRID:SCR_019511, Agilent Technologies, 1260 Infinity, Agilent ZORBAX Eclipse Plus C18 column) to quantify the total amounts of the following amino acids: alanine, glutamate, glutamine, taurine and GABA as previously described (Andersen, Westi, et al., 2021). Briefly, samples were pre-column derivatized with o-phthalaldehyde and fluorescent detection at λex = 338 nm, λem = 390 nm and were eluted using a gradient of increasing concentrations of organic solvent mobile phase (acetonitrile 45%: methanol 45%: H_2_O 10%, v:v:v) and a phosphate-based mobile phase. The amino acid amounts were quantified using calibration curves of external standards of the amino acids of interest with known increasing concentrations. Amounts are expressed as nmol/mg of total cellular protein quantified with the BCA assay kit (Thermo Fisher Scientific Cat. 23225). Selected amino acids for each data set based on their key metabolic and signaling roles in the respective cell types are presented. For microglia, glutamate, glutamine, taurine, and GABA (when detected) were included due to their involvement in microglial metabolism, neuroinflammation, and neurotransmitter modulation. For neurons, aspartate, glutamate, glutamine, and GABA were chosen as they are central to neuronal signaling and the glutamate-glutamine cycle.

### Lactate determination

Lactate released to the incubation media was determined by a L-lactic acid kit (Boehringer Mannheim/R-Biopharm/Roche). Briefly, 50 µL of the incubation media was pipetted into a black 96-well plate, along with L-lactate standard solutions in the range of 1 to 200 µM. Directly after, 105 µL of solution 1+2+3 (1: glycylglycine buffer – pH=10, 2: NAD lyophilizate dissolved in 6 mL H_2_O, 3: Glutamate-pyruvate transaminase suspension) was added to each well, and the fluorescence was read at λ_ex_=350 nm and λ_em_=455 nm, as a background measurement. Subsequently, 50 µL of solution 4 (Lactate dehydrogenase solution) was added to each well and the plate was incubated for 90 minutes at 37°C. Following incubation, the plate was read under the same conditions as outlined earlier. Results were subtracted from background and total lactate content was calculated by constructing a standard curve based on a quadratic equation.

### Immunocytochemistry (ICC)

Following 90 minutes incubation of primary cultures of microglia with the desired substrates, with or without 100 ng/mL LPS, the cells were fixed in 4% PFA for 10 minutes at room temperature (RT), after which they were washed three times with PBS. After fixation, the cells were preincubated with 300 µL permeabilization solution (PBS -/-, 0.2% TritonX-100 and 0.5% Donkey serum) for one hour at 4°C. Subsequently, the cells were incubated overnight at 4°C with 300 µL primary antibody diluted in permeabilization solution. Rabbit anti-IBA1 was diluted 1:500 and chicken anti-GFAP was diluted 1:1000. The following day the primary antibodies were removed, and the cells were washed twice with PBS -/-. After which the cells were incubated with 300 µL secondary antibody diluted (1:200) in permeabilization solution for two hours in darkness. The cells were washed with PBS -/- three times and incubated with DAPI diluted 1:1000 in PBS -/- for 10 minutes in darkness at RT. The cells were washed three times with PBS -/- and subsequently kept in PBS at 4°C until analyzed. Total quantification and classification of microglia by morphology was performed as outlined in He et al. (Y. He et al., 2021). In addition, quantification was carried out in at least three technical replicates per condition with at least five different sections of the wells per treatment. Images were captured using a Leica DM IL LED Fluor Inverted Microscope and images were analyzed using LASX software (RRID:SCR_013673). All images displayed are 40X magnification.

### Statistical analysis

All statistical analyses were performed in GraphPad Prism version 10.1.1 (GraphPad Software Inc., California, USA). The data is depicted as mean ± the standard error of the mean (SEM), with individual data points displayed. In the figure legends, the biological replicates (number of different experimental batches) are denoted “N”, while technical replicates (number of wells for each type of cell culture condition) are denoted “n”. No exclusion criteria were pre-determined; however, one control sample from the [U-^13^C]glutamine incubations was lost during GC–MS analysis. Furthermore, an analytical error resulted in the loss of glutamine M+3 from a LPS sample from the [U^13^-C]GABA incubation. No test for outliers was performed. Data were first tested for normality using either D’Agostino & Pearson or Shapiro–Wilk test (α=0.05). Unpaired data sets were then compared using either Welch’s t-test or Mann–Whitney test, One-way ANOVA was employed for comparing more than two different conditions. The level of significance was set at p < 0.05, denoted by asterisks (*) in the graphs as follows: *p < 0.05, **p <0.01, ***p <0.001 ****p <0.0001. For further details refer to the supplementary statistical report (**SUPP1**).

## Results

### Characterization of microglia cell cultures

Primary microglia cell cultures were validated utilizing immunocytochemistry (ICC). Cell cultures were stained for the microglia specific protein, ionized calcium binding adaptor molecule 1 (IBA1) (Imai et al., 1996; Ito et al., 1998) and glial fibrillary acidic protein (GFAP), a specific marker of astrocytes. Cell nuclei were stained with DAPI. The main cell population in the primary cell cultures were IBA1 positive (**Fig. 1A**), with only minor astrocytic (**GFAP-expressing cells**) contamination. Pro-inflammatory activation following LPS stimulation in primary cultures of microglia has been validated in previous studies (Yingbo He et al., 2021; Horvath et al., 2008; Lund et al., 2006) and was further confirmed in our cultures. Upon LPS activation, the primary microglial cultures exhibit increased expression of pro-inflammatory cytokines (**Fig. 1B**) and undergo glycolysis-predominant metabolic reprogramming accompanied by decreased mitochondrial oxygen consumption (**Fig. 1C**).

**Figure 1:**
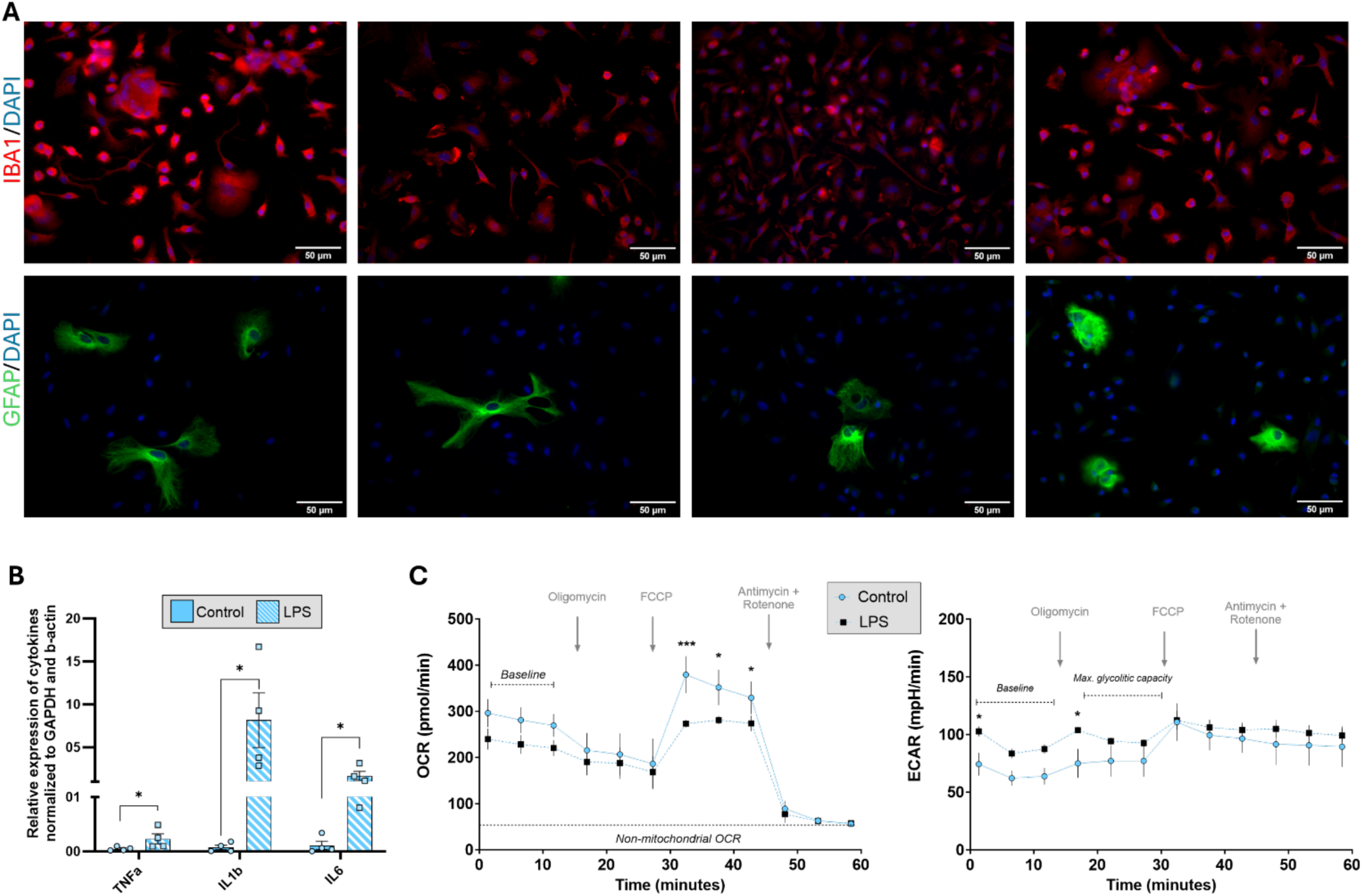
Validation of primary cultures of microglia and characterization of pro-inflammatory activation and metabolic reprogramming. **A.** Representative immunocytochemistry (ICC) images confirming the identity of primary microglia cultures and exhibiting low astrocytic contamination. Cultures were stained with nuclei marker DAPI (blue), IBA1 (red, top panels), specific microglia marker and GFAP (green, lower panels), a specific marker for astrocytes. Representative image of cultures IBA1, GFPA stainings were performed in parallel in samples from the same cell culture from 4 independent cell preparations. **B.** Microglial cultures were treated with either control medium or LPS (100 ng/mL) for 24 h. qPCR analysis showed significantly increased mRNA expression of pro-inflammatory cytokines TNF-α, IL-1β, and IL-6 in LPS-stimulated microglia compared to controls. Mean ± SEM, N (independent cell preparation)=4, n(technical replicates)=1, unpaired t-test, *<0.05. C) Oxygen consumption rate (OCR), reflecting mitochondrial respiration, was reduced, while **C.** extracellular acidification rate (ECAR), indicative of glycolytic activity, was elevated in LPS-treated microglia, as measured using the Seahorse XFe96 analyzer, supporting a shift toward glycolytic metabolism upon activation. Mean ± SEM, N (independent cell preparation)=4, n (technical replicates)=48 wells, two-way ANOVA, *<0.05

### Glycolysis and oxidative glucose metabolism in microglia

Energy metabolism of primary cultures of microglia was investigated under quiescent and LPS-stimulated conditions using isotope tracing with ^13^C labeled metabolic substrates. Glucose is the primary energy substrate of the brain, and metabolism of [U-^13^C]glucose provides an overview of the general metabolic function (Bernier, York, & MacVicar, 2020). Following uptake, [U-^13^C]glucose is metabolized through glycolysis to yield pyruvate M+3, which can be transaminated to alanine M+3 by alanine amino transferase (ALT) or reduced to lactate M+3 by lactate dehydrogenase (LDH) (**Fig. 2**). The ^13^C enrichment of alanine M+3 was found to be slightly decreased in the LPS-stimulated microglia, when compared to quiescent control cells. No significant changes were observed in intracellular lactate M+3 between LPS-stimulated microglia and control, likewise the amount of lactate released to the media from the LPS-stimulated microglia was also unchanged, in contrast to what has previously been reported (Gimeno-Bayón et al., 2014; Voloboueva et al., 2013). Collectively, these results suggest that the extent of glycolytic function may be slightly increased but is overall comparable between quiescent and LPS-stimulated microglia under these conditions.

**Figure 2:**
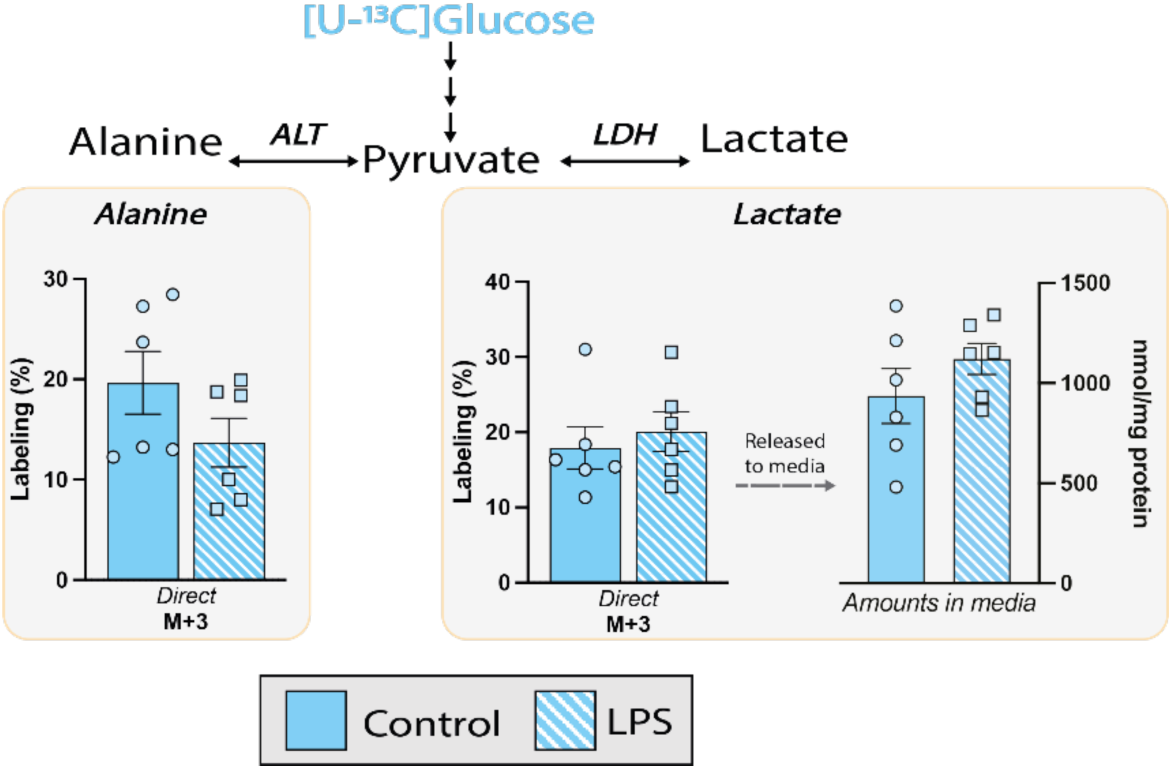
Glycolytic activity in primary cultures of microglia. Intracellular ^13^C enrichments of alanine and lactate and lactate amounts released to media of primary cultures of microglia following metabolism of [U-^13^C]glucose. [U-^13^C]glucose is metabolized via glycolysis resulting in M+3 labeling of the glycolytic end-product pyruvate. As pyruvate is in equilibrium with alanine and lactate, through the enzymes alanine aminotransferase (ALT) and lactate dehydrogenase (LDH), respectively, the ^13^C enrichment in alanine and lactate reflects the glycolytic activity. Mean ± SEM, N (independent cell preparation)=3, n (technical replicates)=1-3, unpaired *t*-test, *<0.05

In the mitochondria, pyruvate M+3 can be converted into acetylCoA M+2 by pyruvate dehydrogenase (PDH) and enter the TCA cycle (**Fig. 3**). AcetylCoA M+2 will give rise to ^13^C enrichment in all TCA cycle intermediates and connected amino acids (M+2 in a first turn and M+3/M+4 in a second turn of the TCA cycle) (A. B. Walls et al., 2014). The ^13^C enrichment from metabolism of [U-^13^C]glucose was overall comparable between LPS-stimulated microglia and control, observed as similar M+2 labeling in citrate, aspartate, glutamate, glutamine and GABA following first turn metabolism of [U-^13^C]glucose (**Fig. 3**). However, enrichment of succinate M+2 was increased in the LPS-stimulated microglia. The unaltered citrate M+2 labeling indicates that the flux of acetylCoA derived from PDH activity was similar between LPS-stimulated microglia and control. Following second turn metabolism of [U-^13^C]glucose, no changes were observed in any of the measured metabolites between LPS-stimulated microglia and control. The overall similar ^13^C enrichment from [U-^13^C]glucose metabolism suggests similar capacity of glucose oxidation in both LPS-stimulated microglia and control.

**Figure 3:**
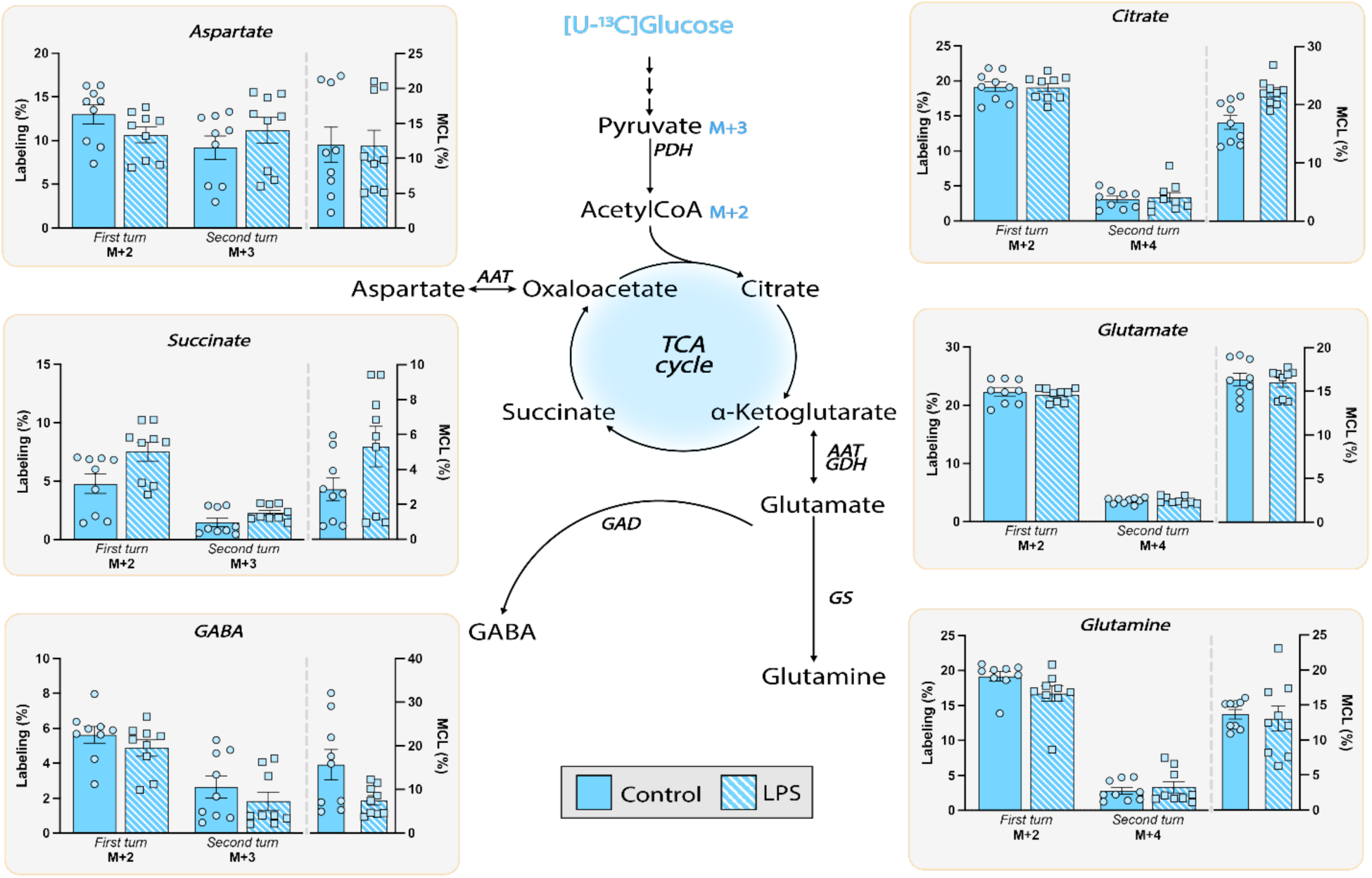
Oxidative metabolism of [U-^13^C]glucose in primary cultures of microglia. Intracellular ^13^C enrichment of TCA cycle intermediates and connected amino acids in primary cultures of microglia following metabolism of 2.5 mM [U-^13^C]glucose. Metabolism of [U-^13^C]glucose occurs through glycolysis to yield pyruvate M+3, which can enter the TCA cycle as acetylCoA M+2, giving rise to ^13^C enrichment in both TCA cycle intermediates and connected amino acids. First turn metabolism of [U-^13^C]glucose is reflected as M+2 labeling, whereas second turn metabolism of [U-^13^C]glucose is represented as M+3/M+4 labeling. Molecular carbon labeling (MCL), calculated from the ^13^C enrichments, reflects the overall accumulation of ^13^C in a given metabolite. AAT: aspartate aminotransferase, GAD: glutamate decarboxylase, GDH: glutamate dehydrogenase, GS: glutamine synthetase, PDH: pyruvate dehydrogenase. Mean ± SEM, N (independent cell preparation)=3, n (technical replicates)=3, unpaired *t*-test with, *<0.05.

The molecular carbon labeling (MCL), being the weighted average percentage of all the isotopologues of a metabolite, provides a measurement of the overall ^13^C accumulation into the TCA cycle (Andersen, Christensen, et al., 2021). Our results showed an overall similar MCL, i.e. ^13^C accumulation in the TCA cycle, between LPS-stimulated microglia and control. However, citrate and GABA showed differences in the MCL, with the latter being decreased, and the former being increased in LPS-stimulated. Collectively, these results suggest that glucose metabolism between LPS-stimulated microglia and control are similar, albeit with a tendency towards higher oxidative capacity in LPS-stimulated microglia when compared to control. Only minor deviations in intracellular amino acids amounts were observed under these conditions where LPS induced a slight but significant increase in GABA intracellular amounts (**Supp. Table 1**).

**Table 1:**
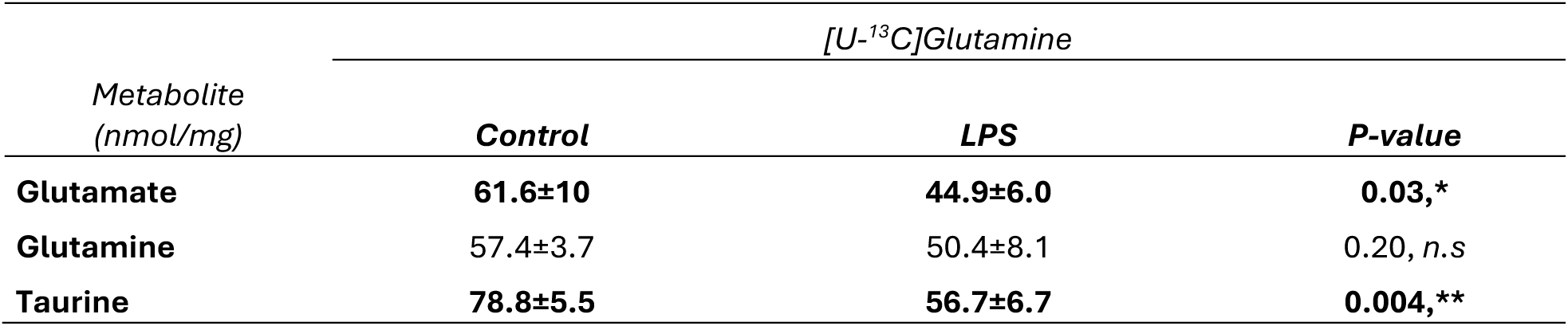
Intracellular amino acid amounts of primary cultures of microglia incubated with [U-^13^C]glutamine. Intracellular amino acid amounts of primary cultures of microglia following incubation with 0.5 mM [U-^13^C]glutamine determined by high performance liquid chromatography (HPLC). Mean ± SEM, N=2, n=3, unpaired *t*-test, *<0.05.

| Metabolite<br>(nmol/mg) | [U-<sup>13</sup>C]Glutamine |  |  |
| --- | --- | --- | --- |
|  | Control | LPS | *P*-value |
| Glutamate | 61.6±10 | 44.9±6.0 | 0.03,* |
| Glutamine | 57.4±3.7 | 50.4±8.1 | 0.20, *n.s* |
| Taurine | 78.8±5.5 | 56.7±6.7 | 0.004,** |

### Uptake and metabolism of glutamine in microglia

Glutamine is synthesized by the astrocyte specific enzyme, glutamine synthetase (GS), and is released into the extracellular space to support neuronal glutamate and GABA synthesis (Andersen & Schousboe, 2023). However, microglia express the glutamine transporter SNAT1 (Jin et al., 2015; Yamada et al., 2019) and possess the necessary cellular machinery to metabolize glutamine (Bernier et al., 2020).

The intracellular labeling of glutamine M+5 was similar between LPS-stimulated microglia and control (Fig. 4). Likewise, the total amounts of intracellular glutamine were similar between LPS-stimulated microglia and control (control vs. LPS: 57.4 ± 0.9 nmol/mg vs. 50.4 ± 1.9 nmol/mg), suggesting similar uptake capacity of glutamine in both conditions (**Table 1**). Intracellularly, glutamine will be converted to glutamate, a reaction catalyzed by the enzyme phosphate-activated glutaminase (PAG). Thus, PAG will convert [U-^13^C]glutamine into glutamate M+5, which can subsequently enter the TCA cycle as α-ketoglutarate M+5. This process results in M+5/M+4 labeling through direct metabolism and M+3/M+2 labeling following first turn metabolism of [U-^13^C]glutamine (**Fig. 4**). The ^13^C enrichment of citrate, succinate, malate and glutamate were all increased in LPS-stimulated microglia compared to control, following direct metabolism of [U- ^13^C]glutamine, whereas no changes were observed in M+5/M+4 labeling of α-ketoglutarate, aspartate and GABA. Following a second turn of the TCA cycle, no changes were observed in the remaining metabolites. Collectively, these results indicate that metabolism of glutamine was increased in LPS-stimulated microglia, when compared to control. Similarly to the incubations with [U-^13^C]glucose, the total amounts of lactate released to the media remained unchanged between the conditions. Furthermore, the total amino acid content was found to be decreased in the LPS-stimulated microglia: glutamate (control vs. LPS: 61.6 ± 2.5 nmol/mg vs. 44.9 ± 1.4 nmol/mg) and taurine (control vs. LPS: 72.4 ± 0.6 nmol/mg vs. 50.9 ± 1.2 nmol/mg) (**Table 1**).

**Figure 4:**
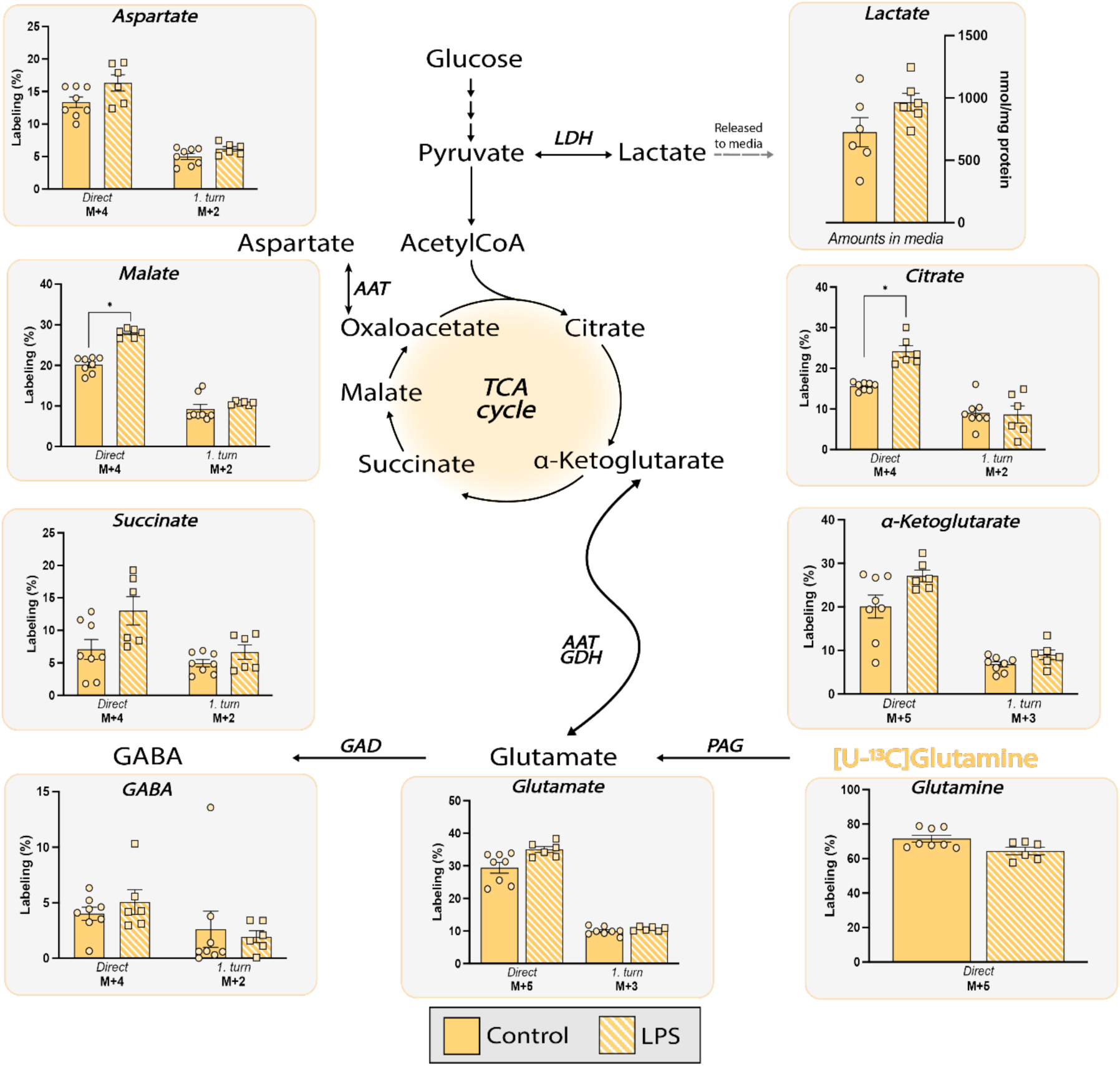
Metabolism of [U-^13^C]glutamine in in primary cultures of microglia. Intracellular ^13^C enrichment of TCA cycle intermediates and connected amino acids in primary cultures of microglia following metabolism of 0.5 mM [U-^13^C]glutamine. Subsequent to uptake, [U-^13^C]glutamine can be converted to glutamate M+5 by phosphate-activated glutaminase (PAG) and enter cellular metabolism as α-ketoglutarate M+5. Direct and first turn metabolism of [U-^13^C]glutamine is reflected as M+5/M+4 and M+3/M+2 labeling, respectively. AAT: aspartate aminotransferase, GAD: glutamate decarboxylase, GDH: glutamate dehydrogenase, LDH: lactate dehydrogenase, PAG: phosphate-activated glutaminase. Mean ± SEM, N (independent cell preparation)=3, n (technical replicates)=2-3, unpaired *t*-test, *<0.05

### Oxidative metabolism of GABA in microglia

GABA is the main inhibitory neurotransmitter and following synaptic transmission it will be taken up by neurons or astrocytes (Zhou & Danbolt, 2013). In astrocytes GABA must undergo oxidative metabolism and a recent study demonstrated that astrocytes may play a more prominent role in GABA metabolism than previously presumed (Andersen et al., 2020). As microglia express GABA transporters in both cell processes and soma they are also capable of neurotransmitter uptake and potentially metabolism (Fattorini et al., 2020). For the carbon backbone of [U-^13^C]GABA to enter cellular metabolism, it must first be converted into succinic semialdehyde by the enzyme GABA-transaminase (GABA-T), subsequently entering the TCA cycle as succinate via the enzyme succinic semialdehyde dehydrogenase (SSADH) (Andersen, Schousboe, & Wellendorph, 2023). Both control and LPS-stimulated microglia were able to efficiently take up [U-^13^C]GABA, leading to 90-95% M+4 enrichment in GABA in both control and LPS-stimulated microglia (**Fig. 5**). However, the total amounts of intracellular GABA were decreased in the LPS-stimulated microglia, when compared to control (control vs. LPS: 20.7 ± 1.0 nmol/mg vs. 14.1 ± 0.2 nmol/mg, respectively), which may indicate decreased GABA uptake capacity in activated microglia (**Table 2**). Direct metabolism of [U-^13^C]GABA in the TCA cycle will give rise to M+4/M+3 enrichment in the TCA cycle and connected amino acids. No changes were observed in succinate M+4, aspartate M+4 and citrate M+4, between control and LPS-stimulated microglia. However, malate M+4, glutamate M+3 and glutamine M+3 were all decreased in the LPS-stimulated microglia when compared to control. Collectively, these findings highlight that microglia are indeed able to take up and metabolize GABA. Furthermore, the results suggest that metabolism between LPS-stimulated microglia and control are similar, albeit with a tendency towards lower metabolism of GABA in LPS-stimulated microglia when compared to control. The total lactate amounts released from microglia to the media in the presence of GABA were similar between control and LPS-stimulated microglia. Interestingly, lower amounts of GABA and taurine were detected in LPS-stimulated microglia compared to control microglia, while no changes were observed in the amounts of glutamate and glutamine (**Table 2**).

**Figure 5:**
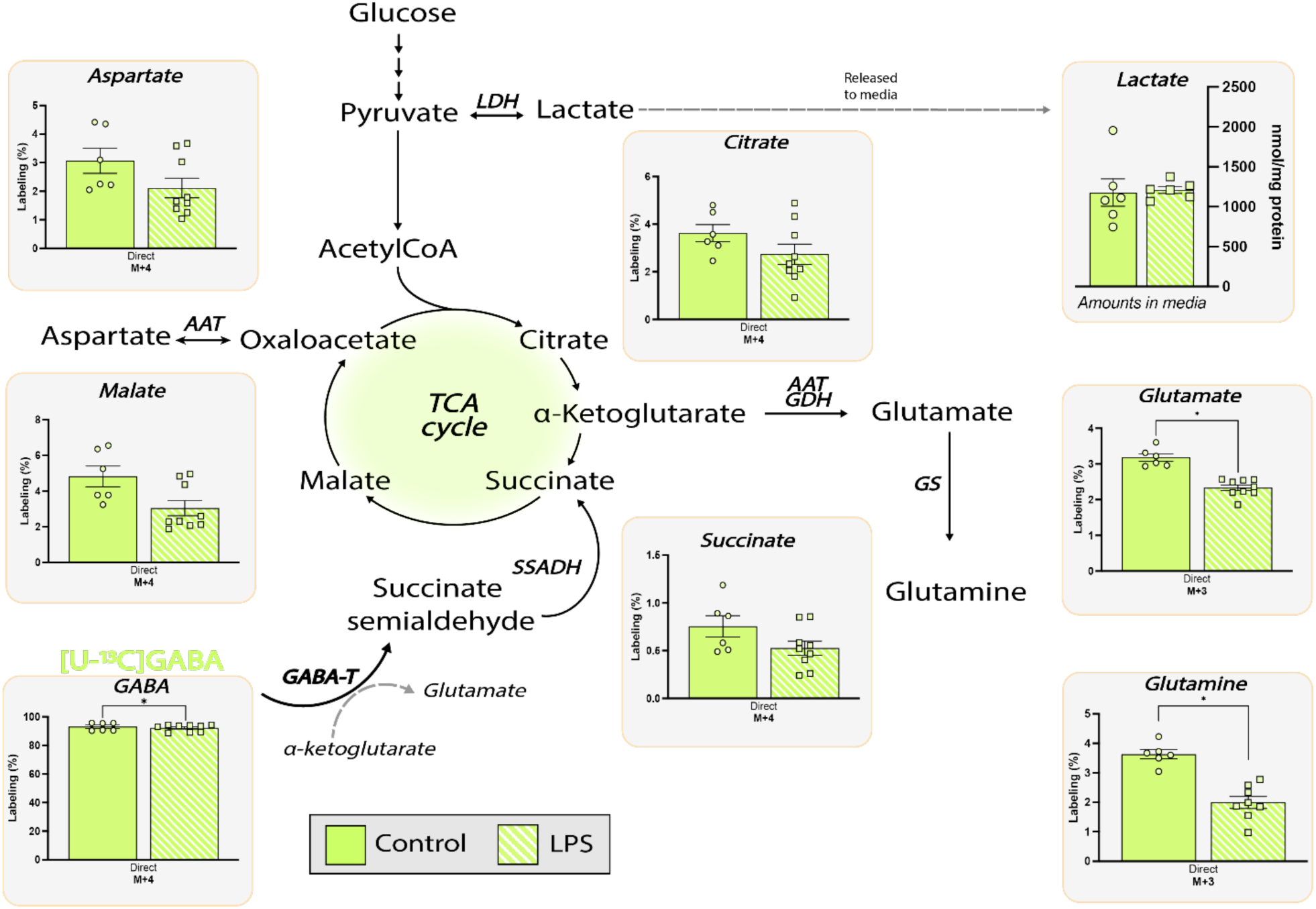
Metabolism of [U-^13^C]GABA acid in primary cultures of microglia. Intracellular ^13^C enrichment of TCA cycle intermediates and connected amino acids in primary cultures of microglia following metabolism of 200 µM [U-^13^C]GABA. Following uptake, [U-^13^C]GABA can be converted to succinate semialdehyde by the enzyme GABA transaminase (GABA-T) and subsequently succinate. Direct metabolism of [U-^13^C]GABA is reflected as M+4/M+3 labeling in TCA cycle intermediates and amino acids. AAT: aspartate aminotransferase, GDH: glutamate dehydrogenase, GS: glutamine synthetase, LDH, lactate dehydrogenase, SSADH, succinic semialdehyde dehydrogenase. Mean ± SEM, N (independent cell preparation)=2-3, n (technical replicates)=3, unpaired *t*-test, *<0.05.

**Table 2:** Intracellular amino acid amounts of primary cultures of microglia incubated with [U-^13^C]GABA. Intracellular amino acid amounts of primary cultures of microglia following incubation with 200 µM [U-^13^C]GABA determined by high performance liquid chromatography (HPLC). Mean ± SEM, N=4, n=3, unpaired *t*-test, *<0.05.

| Metabolite<br>(nmol/mg) | [U-<sup>13</sup>C]GABA |  |  |
| --- | --- | --- | --- |
|  | Control | LPS | P-value |
| Glutamate | 39.5±11 | 33.6±1.9 | 0.68, *n.s* |
| Glutamine | 29.7±9.0 | 22.5±5.1 | 0.46, *n.s* |
| Taurine | **84.8±24** | **51.7±2.4** | **<0.0001,\*\*\*** |
| GABA | **20.7±2.1** | **14.1±1.8** | **0.02,\*** |

### Glucose metabolism in neurons in the presence of quiescent microglia

Having characterized the metabolic flexibility of microglia, we next sought to investigate whether microglia influence neuronal metabolism directly. With this aim, primary cultures of neurons were co-cultured with microglia in a non-contact (NC) system (**Fig. 6A**), and neuronal metabolism was mapped using [U-^13^C]glucose. Monoculture of neurons served as the control group.

**Figure 6:**
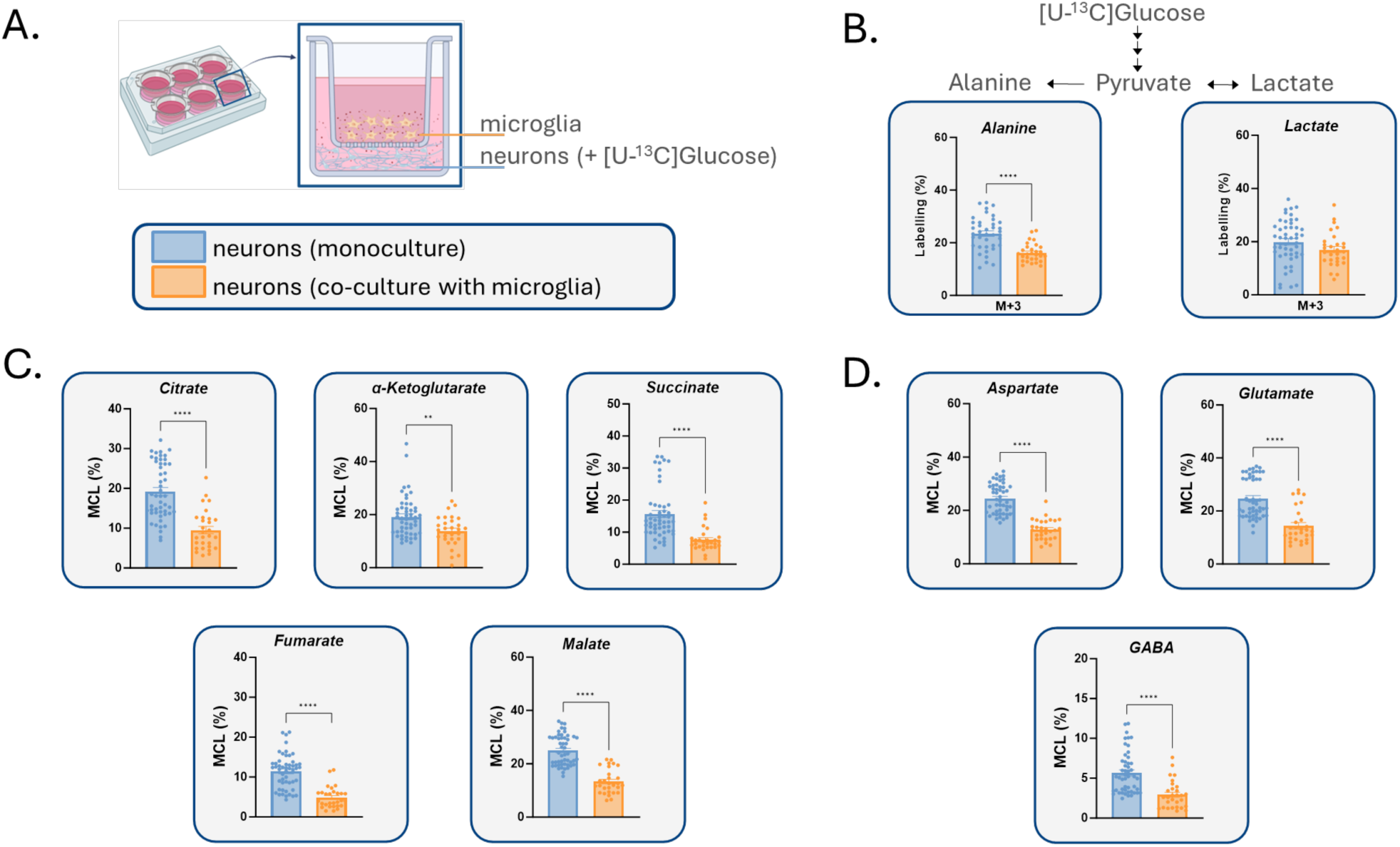
The presence of microglia reduces oxidative metabolism of [U-^13^C]glucose in primary cultures of neurons. **A.** Graphical depiction of the non-contact (NC) co-culture with microglia seeded on filters above neurons. Data from neurons in the NC co-cultures were compared to those in monoculture. Neurons were co-cultured with and without microglia for two hours, with 2.5 mM [U-^13^C]glucose added to the neuronal cultures at the beginning of the second hour. The intracellular ^13^C enrichment in **B.** glycolytic products (alanine and lactate M+3), in **C.** TCA cycle intermediates and **D.** connected amino acids (MCL) in primary cultures of neurons following metabolism of [U-^13^C]glucose. The overall ^13^C-labeling patterns are described in Fig. 2-3. MCL, molecular carbon labeling or the weighted average percentage of all the isotopologues of a metabolite, provides a measurement of the overall ^13^C accumulation Mean ± SEM, N (independent cell preparation)=4-6, n (technical replicates)=6-12, unpaired *t*-test with, *<0.05, **<0.01, ***<0.001, ****<0.0001.

Following incubation with [U-^13^C]glucose, the ^13^C-enrichment in intracellular alanine (M+3) was significantly lower in the neurons co-cultured with microglia compared to the monoculture (**Fig. 6B**). However, the ^13^C-enrichment in intracellular lactate (M+3) was similar in the co-cultured and monocultured neurons. The MCL was determined for TCA cycle intermediates and a significantly lower ^13^C-enrichment in all metabolites was detected in the co-cultured neurons compared to the monoculture (**Fig. 6C**). This effect was also extended to the ^13^C-enrichment in neurotransmitter amino acids, which was lower in the co-cultured neurons (**Fig. 6D**). Collectively, it appears that microglia have a notable impact on neuronal glucose metabolism by significantly decreasing its utilization through the TCA cycle.

The intracellular contents of glutamate and GABA were similar in the co-cultured or monocultured neurons further supporting a lack of cytotoxic effects of microglia on the neurons. However, the amounts of aspartate and glutamine were significantly higher in the co-cultured neurons compared to the monocultured neurons (**Table 3**).

**Table 3:** Intracellular amino acid amounts of primary cultures of neurons incubated with [U-^13^C]glucose in monoculture or NC co-culture with microglia. Intracellular amino acid amounts of primary cultures of neurons following incubation with 2.5 mM [U-^13^C]glucose determined by high performance liquid chromatography (HPLC). Mean ± SEM, N=3-5, n=1-3, unpaired *t*-test, *<0.05.

| Metabolite<br>(nmol/mg) | Glucose |  |  |
| --- | --- | --- | --- |
|  | Monoculture | Co-culture | P-value |
| **Aspartate** | **58.4±8.1** | **100.6±29** | **0.04, \*** |
| **Glutamate** | 113.0±10 | 89.5±14 | 0.14, *n.s.* |
| **Glutamine** | **5.6±0.7** | **7.7±0.9** | **0.03,\*** |
| **GABA** | 47.9±4.4 | 52.4±14 | 0.67, *n.s.* |

### Glucose metabolism in neurons in the presence of activated microglia

Activated microglia undergo significant metabolic reprogramming, favoring glycolytic flux and reducing mitochondrial oxidative metabolism (Aldana, 2019). Given that stimuli like LPS and Aβ (causally associated with Alzheimer’s disease) can induce a pro-inflammatory state in microglia, and our findings suggest that pro-inflammatory stimuli (e.g., LPS) affect microglial metabolism, we investigated whether activated microglia influence neuronal metabolism.

Microglia cultures were stimulated with either LPS or Aβ_1-42_ for 24 hours before being introduced into the co-culture with neurons. Neurons were then incubated with media containing [U-^13^C]glucose in the presence of the activated microglia but in absence of the stimuli (**Fig. 7A** method overview). ^13^C-Enrichment in glycolytic products (alanine M+3 and lactate M+3) showed no significant difference between neurons co-cultured with quiescent or LPS-stimulated microglia (**Fig. 7B**). However, the incorporation of ^13^C in alanine and lactate was higher in neurons co-cultured with Aβ-stimulated microglia compared to neurons co-cultured with quiescent microglia, yet only reaching significance for alanine. The ^13^C-enrichment in neuronal TCA cycle metabolites (MCL) revealed that the LPS-stimulated microglia mirrored the effect of non-stimulated microglia, except for a significant increase in α-ketoglutarate and fumarate labeling. Aβ-stimulated microglia induced a significantly higher ^13^C-enrichment in neuronal citrate and succinate and induced a trend towards higher enrichment in α-ketoglutarate and malate compared to neurons co-cultured with non-stimulated microglia. ^13^C-

**Figure 7:**
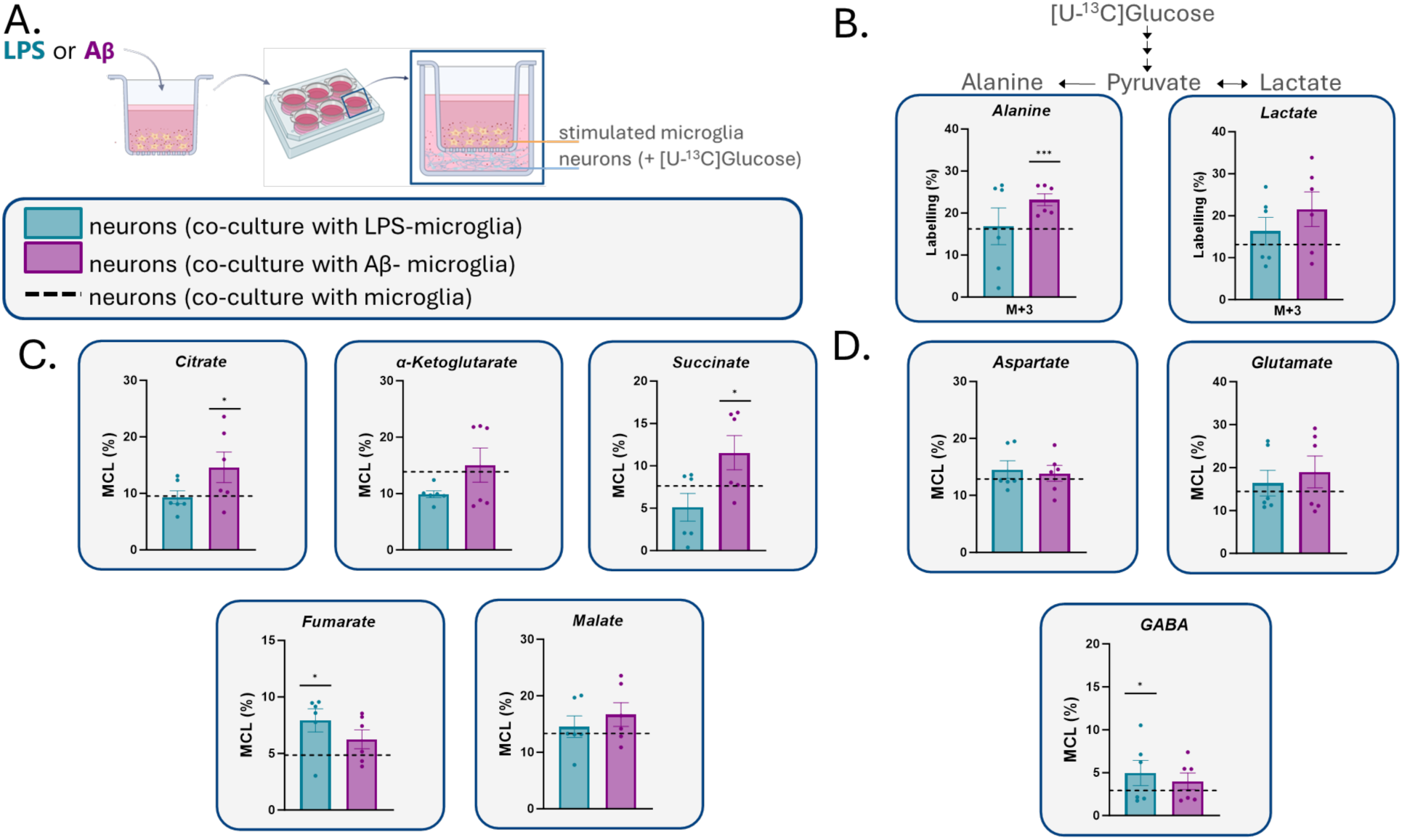
The presence of activated microglia differentially influences neuronal glucose metabolism depending on the activating stimuli. **A.** Graphical depiction of microglia activation and non-contact (NC) co-culture with microglia seeded on filters above neurons. Microglia activation was induced with either LPS or Aβ1-42 for 24h before co-culturing them with neurons. Data from neurons in the NC co-cultures with activated microglia were compared to neurons in NC co-cultures with quiescent microglia (dotted line). Neurons were co-cultured with microglia for two hours, with 2.5 mM [U-^13^C]glucose added to the neuronal cultures at the beginning of the second hour. The intracellular ^13^C enrichment in **B.** glycolytic products (alanine and lactate M+3), in **C.** TCA cycle intermediates and **D.** connected amino acids (MCL) in primary cultures of neurons following metabolism of [U-^13^C]glucose. MCL, molecular carbon labeling or the weighted average percentage of all the isotopologues of a metabolite, provides a measurement of the overall ^13^C accumulation. Mean ± SEM, N (independent cell preparation)=3, n (technical replicates)=2, unpaired *t*-test with, *<0.05 comparing ^13^C-enrichment from neurons co-cultured with quiescent microglia (dotted line), and LPS/Aβ-stimulated microglia.

Enrichment in aspartate and glutamate was unchanged in the neurons co-cultured with LPS- or Aβ-stimulated microglia compared co-cultured neurons with non-stimulated microglia. Only the ^13^C-enrichment in GABA was higher in the neurons co-cultured with LPS-stimulated microglia compared to the non-stimulated microglia (**Fig. 7C-D**).

Additionally, the intracellular amino acid amounts from the neurons co-cultured with LPS- or Aβ-stimulated microglia were compared to the content from the neurons co-cultured with unstimulated microglia. The intracellular content of aspartate was significantly lower in the neurons co-cultured with LPS-stimulated microglia compared to the neurons co-cultured with quiescent microglia, while the amounts of glutamate, glutamine and GABA were similar between all co-cultured neurons (**Table 4**). These results indicate that LPS- and Aβ-stimulated microglia may differentially affect neuronal metabolism.

**Table 4:** Intracellular amino acid amounts of primary cultures of neurons incubated with [U-^13^C]glucose in NC co-culture with stimulated microglia. Intracellular amino acid amounts of primary cultures of neurons following incubation with 2.5 mM [U-^13^C]glucose determined by high performance liquid chromatography (HPLC). Microglia were stimulated with LPS or Aβ1-42 for 24h, after which time, the stimuli were removed and the microglia cultures were put in contact with neurons. Mean ± SEM, N=3, n=2, **^1.^** unpaired *t*-test, *<0.05 comparing ^13^C-enrichment from neurons co-cultured with quiescent microglia (values in table 3), and LPS/Aβ-stimulated microglia.

| Metabolite<br>(nmol/mg) | Glucose |  |  |  |
| --- | --- | --- | --- | --- |
| | Co-culture (neurons and<br>LPS- microglia) | P-value<sup>1</sup> | Co-culture<br>(neurons and Aβ-<br>microglia) | P-value<sup>1</sup> |
| Aspartate | 45.9±10 | 0.047, * | 85.9±20 | 0.58, *n.s.* |
| Glutamate | 89.2±26 | 0.99, *n.s.* | 108.5±14 | 0.23, *n.s.* |
| Glutamine | 7.9±2.5 | 0.85, *n.s.* | 6.7±1.5 | 0.46, *n.s.* |
| GABA | 31.6±11 | 0.13, *n.s.* | 55.0±6.3 | 0.81, *n.s.* |

To exclude the possibility that the LPS-dampening effect of microglia on neuronal metabolism was due to microglia-induced neuronal death, we performed a cell viability assay on neurons in the presence and absence of quiescent and LPS-stimulated microglia. Compared to the neurons in monoculture, we did not detect significant changes in cell viability in the neurons co-cultured with quiescent nor with LPS-stimulated microglia. (**Supp. Fig. 1**).

## Discussion

Our study provides new insights into the metabolic flexibility of microglia and their impact on neuronal metabolism, highlighting the intricate interplay between immune and neuronal cells in the CNS.

Immunometabolism is a relatively new field that emerged upon the discovery that peripheral macrophages were able to shift metabolic phenotype (Bernier et al., 2020; Viola et al., 2019; York et al., 2021). As the resident immune cell of the brain, microglia share cellular characteristics with peripheral macrophages. Recent studies have sought to elucidate microglia metabolism utilizing indirect measurements of cellular metabolism such as two-photon imaging or live-cell bioenergetics (Bernier, York, & MacVicar, 2020). Here, we demonstrate that microglia are metabolically flexible cells capable of utilizing different energy substrates, as has been reported in macrophages. In addition, we found that activated microglia change their metabolic phenotype and exerts specific effect on neuronal metabolism depending on the activation stimuli. Notably, we show for the first time that microglia are able to take up and metabolize GABA, which has not previously been suggested or investigated.

Our findings show that LPS stimulation does not significantly modify the ^13^C-enrichment in intracellular lactate, or the amounts of lactate released into the media from our microglia cultures. However, we do observe an increased lactate-derived extracellular acidification upon LPS stimulation. The shift towards glycolysis, often referred to as the Warburg effect, is a hallmark of activated immune cells and has been observed in microglia during pro-inflammatory states (Palsson-McDermott & O’Neill, 2013). As such, our results are puzzling, as these partially conflict with previously published results (Gimeno-Bayón et al., 2014; Nair et al., 2019; Voloboueva et al., 2013; York et al., 2021). One possible explanation for the discrepancies may be how the lactate amounts are measured. The majority of studies reporting increased release of lactate to the extracellular media make use of either the extracellular acidification rate (ECAR) as in our study (Hu et al., 2020; Nair et al., 2019; Voloboueva et al., 2013) or fluorescent lifetime imaging techniques (York et al., 2021), whereas we additionally measure lactate amounts directly in the media. Furthermore, it is important to consider the potential activation of the pentose phosphate pathway (PPP) as a key component of microglial metabolic reprogramming in response to LPS stimulation(Gimeno-Bayón et al., 2014; Tu et al., 2019). Upon TLR4 activation, microglia typically undergo a dual metabolic shift: glycolysis is upregulated to meet immediate bioenergetic demands and support lactate production, while the PPP is activated to generate NADPH, which is essential for reactive oxygen species (ROS) production, glutathione recycling, and overall redox homeostasis (Peruzzotti-Jametti et al., 2021). This interplay enables microglia to sustain prolonged inflammatory responses while managing oxidative stress. In our study, the unchanged lactate levels observed at early time points may reflect a transient phase in which PPP activation precedes or occurs independently of overt glycolytic upregulation. Increased PPP flux could be driven by early glutathione turnover and NADPH demand, aligning with previous studies showing that PPP activation can serve as an initial immunometabolic adaptation prior to broader shifts in glycolysis (Huang et al., 2024). Further investigation using PPP-specific tracers and direct NADPH quantification will be valuable for elucidating these early-phase metabolic dynamics and their implications for microglial function and neuroinflammation.

Interestingly, the oxidative metabolism of glucose, as indicated by the ^13^C enrichment in TCA cycle intermediates, remained largely unchanged between LPS-stimulated and quiescent microglia. This might suggest that both glycolysis and the capacity for oxidative phosphorylation are maintained. However, following first turn metabolism of [U-^13^C]glucose, the labeling in succinate M+2 was found to be higher in the LPS-stimulated microglia, when compared to control. In peripheral macrophages stimulated with LPS, intracellular amounts of succinate are increased and functions as a proinflammatory marker that can drive the inflammatory response (Mills et al., 2016; Tannahill et al., 2013). No other changes were observed between the different conditions following both first and second turn metabolism of [U-^13^C]glucose. To corroborate this, we report lower intracellular taurine amounts in LPS-stimulated microglia. As taurine is an amino acid involved in osmoregulation and antioxidant defense, the lower amounts may reflect altered cellular consumption, uptake or release in response to activation and oxidative stress induced by LPS (Park et al., 2002) (Supp. Table 1). Taurine uptake is mediated by the taurine transporter (TauT, SLC6A6), which may be regulated by inflammatory signals. In line with this, it has been reported that LPS reduces taurine uptake in enterocytes (O’Flaherty et al., 1997). Additionally, LPS-induced inflammation is associated with changes in cysteine metabolism in macrophages (Takeda et al., 2023), a precursor for taurine biosynthesis. Increased cysteine demand for glutathione synthesis under oxidative stress could lead to reduced taurine synthesis. Conversely, LPS-stimulated microglia had higher intracellular amounts of glutamine following incubation with [U-^13^C]glucose in LPS-stimulated microglia compared to control (supp. Table 1), which may suggest an increased demand for this amino acid to support the oxidative phosphorylation during inflammation.

Glutamine is a critical substrate for various metabolic pathways, including the TCA cycle (Andersen & Schousboe, 2023; Bradford, Ward, & Thomas, 1978). Recently, it was reported that glutamine was able to sustain *in situ* cellular metabolism in microglia, even in the absence of glucose (Bernier et al., 2020). To further elucidate the extent of glutamine metabolism in microglia, [U-^13^C]glutamine was employed. Glutamine is taken up into microglia via the SNAT1 glutamine transporter (Bernier, York, & MacVicar, 2020), after which it can enter oxidative metabolism through conversion to glutamate and subsequently α-ketoglutarate. We demonstrate that quiescent microglia readily take up and metabolize glutamine. This aligns with previous findings, showing that microglia, *in situ*, are able to sustain cellular functions in aglycemic conditions when metabolizing glutamine (Bernier et al., 2020). However, uptake of glutamine was similar between LPS-stimulated microglia and quiescent microglia, as evident from table 1 and Fig.4. The similar uptake capacities of glutamine suggest that the observed metabolic changes are due to alterations in intracellular metabolism rather than differences in substrate availability. However, direct metabolism of glutamine in LPS-stimulated resulted in higher ^13^C enrichment in glutamate, succinate, malate and citrate. These findings are in line with a recent report showing that LPS-stimulated primary cultures of microglia display a higher capacity for glutamine oxidation (Zhang et al., 2024). Interestingly, when glutamine metabolism was blocked pharmacologically, microglial inflammatory pathways were also inhibited (Zhang et al., 2024). Taken together, this may indicate that glutamine metabolism in microglia is necessary to drive inflammation. LPS-stimulated microglia also had lower intracellular taurine amounts, again suggesting that this amino acid might be either consumed or its uptake or release is altered in response to inflammation.

Of note is the comparable labeling of citrate and succinate from [U-^13^C]glucose and [U-^13^C]glutamine. Although it could indicate equal metabolic contributions of the two substrates, it most likely reflects the dynamic and interconnected nature of TCA cycle metabolism, where carbon atoms from different entry points (acetyl-CoA vs. α-ketoglutarate) are redistributed across intermediates through multiple turns and enzymatic exchanges. Additionally, factors such as metabolite pool sizes, turnover rates, and cellular heterogeneity can further influence isotopic enrichment, which make interpretations of substrate preference based solely on labeling intensity quite complex. The ability of microglia to take up and metabolize GABA, the primary inhibitory neurotransmitter, highlights their metabolic versatility. To our knowledge, this is the first study to address functional GABA metabolism in microglia. We have recently demonstrated that astrocytes are capable of metabolizing GABA to a large extent in both intact brain tissue and cultured cells (Andersen et al., 2020; Salcedo et al., 2021). However, the primary microglia cell cultures are almost void of astrocytic contamination, and therefore the metabolic mapping of [U-^13^C]GABA can only be attributed to microglia. Both LPS-stimulated and quiescent microglia were able to take up [U-^13^C]GABA to a large extent. However, the intracellular amounts of GABA were significantly lower in LPS-stimulated microglia, when compared to control (Table 2), which is in line with what has been observed in macrophages (Stuckey et al., 2005). Subsequent to uptake, [U-^13^C]GABA enters oxidative metabolism giving rise to M+4/M+3 labeling following direct metabolism. The ^13^C enrichment in all TCA cycle intermediates and connected amino acids were all below 5%, suggesting that GABA metabolism is generally low, but active nonetheless in primary microglia. Overall, metabolism of [U-^13^C]GABA was similar between LPS-stimulated and quiescent microglia. Interestingly, lower ^13^C enrichment was observed in both glutamine and glutamate in LPS-stimulated microglia. This is in contrast to what has previously been observed in both primary cultures of activated microglia (Zhang et al., 2024) and macrophages (Stuckey et al., 2005). This may indicate that GABA modulates the production and release of amino acids in activated microglia. Furthermore, the extracellular release of lactate, a marker for glycolytic activity, was comparable between LPS-stimulated and quiescent microglia, suggesting that GABA may help maintain metabolic homeostasis in microglia by stabilizing glycolytic flux. Additionally, this stabilization could be linked to GABA’s role in modulating proinflammatory responses(Kang et al., 2022) (Bhandage et al., 2018; Bhat et al., 2010).

Our co-culture experiments revealed that the presence of microglia, particularly when activated by LPS or Aβ, significantly impacts neuronal glucose metabolism. The decreased ^13^C enrichment in TCA cycle intermediates and neurotransmitter amino acids in neurons co-cultured with microglia suggests a reduction in the neuronal metabolic rate and neurotransmitter metabolism. This could be mediated through several indirect mechanisms, including changes in energy substrate availability, neurotransmitter homeostasis, inflammatory signaling, and oxidative stress. The metabolic demands of microglia may reduce the availability of glucose by competing with the neurons for glucose uptake and utilization. In addition, lactate release from microglia could provide neurons with an auxiliary fuel. Our findings indicate that microglia can take up and metabolize GABA. Thus, if microglia deplete extracellular GABA levels, the inhibitory neuronal tone may be reduced, potentially leading to compensatory reductions in neuronal firing and metabolism. Moreover, microglia are known to produce radical oxygen species (ROS) and nitric oxide, which may impair neuronal metabolism by inhibiting mitochondrial electron transport chain activity. Alternatively, neurons may downregulate metabolic activity as a protective response to mitigating oxidative damage. Future research is advocated to elucidate the mechanisms underlying the metabolic response of neurons to microglia.

Another key finding of this study is that the presence of microglia exerts differential effects on neurons depending on the type of microglia-activating stimulus. This is in line with the observations that LPS and Aβ activate microglia through distinct pathways eliciting differential responses. This may lead to different metabolic patterns in neurons when co-cultured with LPS- or Aβ-stimulated microglia. LPS specifically activates toll-like receptor 4 (TLR4) in microglia, triggering intracellular cascades that result in a robust inflammatory response, characterized by the release of pro-inflammatory cytokines (TNF-α, IL-1β, IL-6), inducible nitric oxide synthase, and increased cytosolic Ca²⁺ concentrations (Hines et al., 2013; Qin et al., 2005). Conversely, Aβ has been shown to interact with both TLR4 and triggering receptor expressed on myeloid cells 2 (TREM2) in microglia (Frank et al., 2008; Walter et al., 2007). TREM2 activation leads to microglial proliferation, lipid metabolism, phagocytosis, and inflammatory responses. However, mild TREM2 activation can inhibit the pro-inflammatory response induced by TLR4, leading to an anti-inflammatory effect (Liu et al., 2020; Ulland et al., 2017). A growing body of evidence indicates that Aβ profoundly alters microglial metabolism, driving these cells toward a glycolytic, pro-inflammatory state that perpetuates neuroinflammation and neuronal injury. This metabolic reprogramming involves not only increased glycolytic flux, but reduced mitochondrial respiration, and changes in key metabolic regulators such as HIF-1α and mTOR (Baik et al., 2019; Lauro & Limatola, 2020; Lu et al., 2021). Such alterations not only support the energetic needs of activated microglia but may also contribute to their chronic activation and impaired phagocytic function. Importantly, these metabolic changes can be further influenced by the cellular environment, including factors secreted in co-culture systems, highlighting the complexity of neuron-glia metabolic interactions. In our co-culture system, it is plausible that the presence of Aβ-stimulated microglia could further alter their own metabolic state, as well as that of neighboring neurons, even in non-contact conditions. Soluble factors released by Aβ-exposed microglia (e.g., cytokines, metabolites) could mediate metabolic crosstalk, amplifying or modifying the metabolic phenotype of both cell types. The reciprocal influence of neurons and glia in such systems is an emerging area of research, and future studies employing will be critical to dissect these interactions.

The increased ^13^C enrichment in alanine, lactate as well as in the TCA cycle intermediates, citrate and succinate, in neurons co-cultured with Aβ-stimulated microglia suggests a compensatory increase in glucose metabolism, likely as a response to microglia activation. These findings are consistent with reports that neuroinflammation can disrupt neuronal energy metabolism, contributing to the pathogenesis of diseases such as AD. Interestingly, the ^13^C-enrichment in GABA was selectively higher in the neurons co-cultured with LPS-stimulated microglia but not with Aβ-stimulated microglia, which suggests differential metabolic effects of these microglial activation states on neuronal GABA metabolism. Specifically, LPS-stimulated microglia could provide metabolic substrates or signals that enhance neuronal GABA synthesis - a response not observed with Aβ-stimulated microglia. LPS-stimulated microglia may release precursors for GABA synthesis including glutamine, which neurons can then take up and convert to GABA. Proinflammatory cytokines (e.g., IL-1β, TNF-α) may modulate neurotransmitter metabolism (Miller et al., 2013), therefore it can be speculated that LPS-induced cytokines may enhance GAD expression or activity, promoting GABA synthesis. In general, higher neuronal GABA synthesis in response to LPS suggests a compensatory mechanism where neurons may upregulate inhibitory neurotransmission to counteract the hyperexcitability associated with neuroinflammation (Kurki et al., 2023).

Failure of Aβ-stimulated microglia to induce similar metabolic changes could contribute to dysfunctional inhibitory signaling in neurodegenerative diseases like in Alzheimer’s disease (AD), where impaired GABAergic function is observed (Bi et al., 2020). A similar metabolic role in relation to GABA homeostasis has been proposed for astrocytes in AD (Andersen, Schousboe, & Verkhratsky, 2022). LPS-stimulated microglia may enhance glutamate-to-GABA conversion, whereas Aβ-stimulated microglia fail to support this metabolic pathway, potentially contributing to neurotransmitter imbalance in neurodegenerative conditions. It has been reported that pro-inflammatory stimulation induces microglia to release glutamate through the cystine/glutamate antiporter (also known as Xc exchange system) (Barger et al., 2007). This antiporter exchanges intracellular glutamate for extracellular cystine, which is essential for glutathione synthesis and antioxidant defense. Upon activation by stimuli such as LPS, microglia upregulate the Xc system, leading to increased glutamate export as part of their response to oxidative stress and heightened demand for cysteine. This mechanism links microglial redox homeostasis directly to extracellular glutamate accumulation, which can contribute to excitotoxic neuronal injury during neuroinflammation. In light of this, our data on microglial glutamate handling likely reflect not only changes in neurotransmitter metabolism but also this stress-adaptive, yet potentially neurotoxic, pathway. It can be speculated that heightened microglial glutamate export may overstimulate neuronal glutamate receptors, especially extrasynaptic NMDA receptors. This excitotoxic signaling may lead to excessive calcium influx, mitochondrial dysfunction, and increased ROS production in neurons, impairing their bioenergetic capacity. As a result, neurons may experience disrupted mitochondrial respiration and reduced ATP generation, manifesting as hypometabolism. Moreover, the excitotoxic environment can trigger synaptic loss and impair neurotransmission, further diminishing neuronal energy demand and function. Thus, microglia-derived glutamate release may create a feed-forward loop where oxidative stress and excitotoxicity converge to suppress neuronal metabolism. Further studies are needed to elucidate these specific mechanisms and to explore whether targeting microglial metabolism can modulate neuronal inhibitory and excitatory signaling and protect against neurodegeneration.

In summary, our study provides comprehensive insights into the metabolic flexibility of microglia and their impact on neuronal metabolism. The upregulation of glutamine metabolism in LPS-stimulated microglia highlights their metabolic adaptability during inflammation. The differential effects of LPS and Aβ-stimulated microglia on neuronal glucose metabolism underscore the complex interplay between neuroinflammation and neuronal function. These findings contribute to our understanding of microglial immunometabolism and its implications for neurodegenerative diseases.

## ABBREVIATION LIST

AAT: Aspartate aminotransferase
AD: Alzheimer’s disease
ALT: Alanine amino transferase
Aβ: Amyloid-β
CNS: Central Nervous system
ECAR: Extracellular acidification rate
GABA: γ-aminobutyric acid
GABA-T: GABA transaminase
GC-MS: Gas chromatography-mass spectrometry
GDH: Glutamate dehydrogenase
GS: Glutamine synthetase
HPLC: High-Performance liquid chromatography
ICC: Immunocytochemistry
LDH: Lactate dehydrogenase
LPS: Lipopolysaccharide
MCL: Molecular carbon labelling
NC: Non-contact (co-culture system)
PAG: Phosphate activated glutaminase
PDH: Pyruvate dehydrogenase
PPP: Pentose phosphate pathway
ROS: Radical oxygen species
RRIDs: Research Resource Identifier (see scicrunch.org)
SSADH: Succinic semialdehyde dehydrogenase
TCA: Tricarboxylic acid (cycle)
TLR4: Toll-like receptor 4
TREM2: Triggering receptor expressed on myeloid cells 2

## Supplementary Materials

The following supporting information complements the study:

SUPP1 Full statistical report

Supp. methods

Supp. Figure 1. The presence of microglia does not affect neuronal survival.

Supp. Table S1: Intracellular amino acid amounts of primary cultures of microglia incubated with 2.5 mM [U-^13^C]glucose.

## Author Contributions

Conceptualization, B.I.A., E.W.W; methodology, E.W.W., R.B. C., Z. K., D. A., B.G.S, A.C.C., J.V.A., N.K.H., S.S.M., C.G., B. I. A.; formal analysis, E.W.W., B.I.A., R.B.C., A.C.C, J.V.A, N.K.H. S.S.M.,; investigation, E.W.W., R.B. C., Z. K., D. A., A.C.C., J.V.A., N.K.H., S.S.M., B. I. A; resources, B.I.A; data curation, E.W.W., R.B. C., A.C.C., J.V.A., B. I. A.; writing—original draft preparation, E.W.W., R.B. C., J.V.A., B. I. A; writing—review and editing, all co-authors; funding acquisition, B.I.A. All authors have read and agreed to the published version of the manuscript.

## Data Availability Statement

Data is provided in the full statistical report (SUPP1) or can be provided upon reasonable request by contacting the authors.

## Acknowledgments

The authors would like to acknowledge the excellent technical assistance of laboratory technician Heidi Marie Nielsen.

## SUPPLEMENTARY MATERIAL

### SUPP1 Full Statistical Report (pdf file)

#### Supp. Methods

##### Quantitative polymerase chain reaction (qPCR)

*RNA Extraction.* Primary microglia were incubated for 24h with metabolic substrates ± 100 ng/mL LPS, washed once with PBS, and lysed in 350 µL RLT buffer. Lysates were mixed with an equal volume of 70% ethanol and processed using RNeasy Mini spin columns (Qiagen) following the manufacturer’s protocol, including on-column DNase I treatment. RNA was eluted in 40 µL RNase-free water and stored at –20°C. *RNA Quantification and cDNA Synthesis.* RNA concentration was measured with a NanoDrop 1000 (RRID:SCR_016517, Thermo Fisher Scientific) and samples were diluted to the lowest concentration of all samples. Due to low RNA yield (<100 ng/µL), cDNA synthesis and whole transcriptome amplification were performed using the QuantiTect Whole Transcriptome Kit (Qiagen, RRID:SCR_008539) according to the manufacturer’s instructions. The final average cDNA concentration was 2350 ± 90.2 ng/µL. *qPCR Amplification.* Primers (10 µM, see below) were prepared for target and reference genes. cDNA was diluted to 50 ng/µL. qPCR reactions contained 10 µL 2X SYBR Green (Roche), 0.8 µL each primer, 6.4 µL RNase-free water, and 2 µL cDNA (final volume 20 µL). Reactions were performed on a MyGo Pro qPCR system (Cork, Ireland) with the following cycling: 95°C for 15 min; 45 cycles of 95°C for 15 s, 58°C for 30 s, 72°C for 30 s; final extension at 72°C for 10 min; followed by melting curve analysis. Gene expression was normalized to GAPDH and analyzed using the 2^−ΔΔCT^ method using GraphPad Prism.

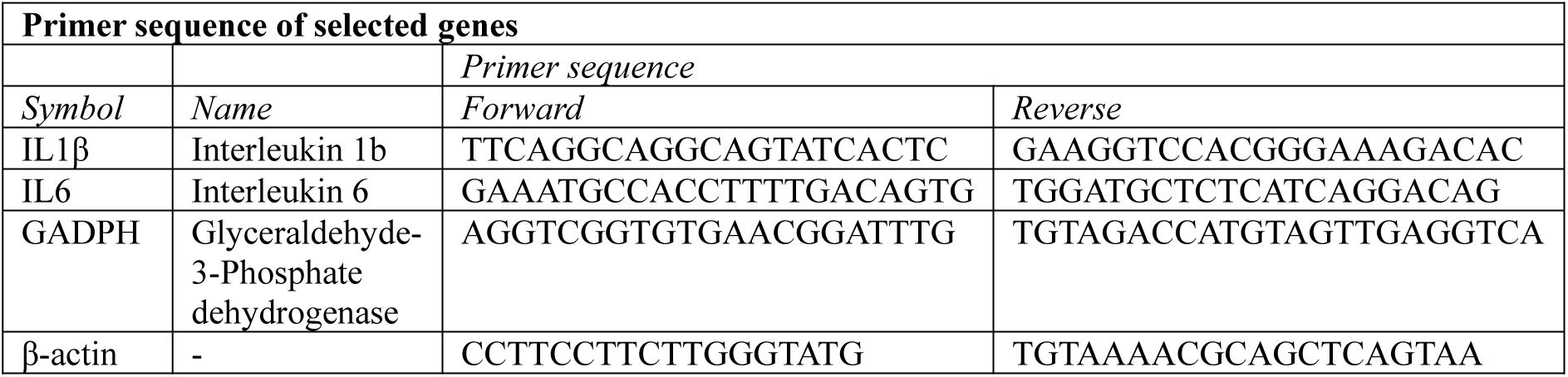

##### Seahorse XFe96 Assay

Real-time oxygen consumption rate (OCR) and extracellular acidification rate (ECAR) were measured using a Seahorse XFe96 analyzer (RRID:SCR_019545, Agilent Technologies, Santa Clara, CA, USA). Primary cultures of microglia cells were seeded in Seahorse micro plates and incubated at 37°C with 5% CO₂. Where indicated, cells were stimulated with 100 ng/mL LPS 24 hours before the assay. Before the assay, the medium was replaced with Seahorse assay medium (DMEM, 2.5 mM glucose, pH 7.4, without Na₂CO₃), and plates were incubated for 60 minutes in a non-CO₂ incubator at 37°C. Baseline OCR and ECAR were recorded over three measurement cycles (1 min mix, 0.5 min wait, 3 min measure). Subsequently, the following compounds were sequentially injected: oligomycin (OM, final 1 µM), FCCP (1 µM), and rotenone/antimycin A (Rot/AA, 0.5 µM each), with three measurement cycles after each injection. Data analysis was performed using Seahorse Wave software (RRID:SCR_014526), which calculated OCR/ECAR along with mitochondrial parameters including basal respiration, proton leak, maximal respiration, spare respiratory capacity, and non-mitochondrial oxygen consumption.

#### Supp. Figures

**Supp Figure 1.**
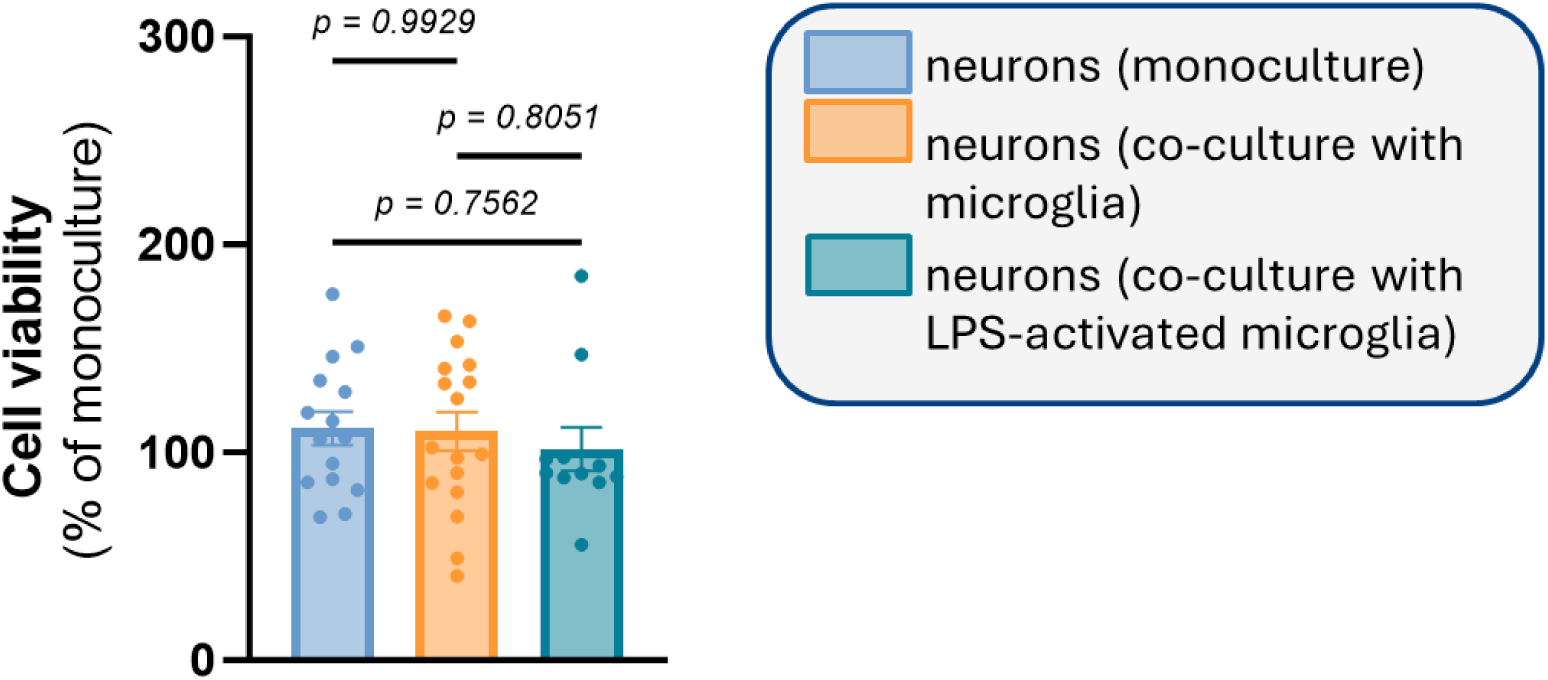
The presence of microglia does not affect neuronal survival. Neurons were co-cultured without microglia (monoculture), or with quiescent or LPS-stimulated microglia for two hours after which time, cell survival was assessed via XTT Assay. Data is normalized with respect to the monoculture analyzed in parallel in each corresponding experimental set (percentage of monoculture). Mean ± SEM, N=3, n=3-6, one-way ANOVA. *P* values are shown, no statistical differences were detected with a threshold of p <0.05 for significance.

**Supp. Table 1.** Intracellular amino acid amounts of primary cultures of microglia incubated with 2.5 mM [U-^13^C]glucose. Intracellular amino acid amounts of primary cultures of microglia following incubation with 2.5 mM [U-^13^C]glucose determined by high performance liquid chromatography (HPLC). Mean ± SEM, N=4, n=3, unpaired *t*-test, *<0.05.

| *Metabolite*<br>(nmol/mg) | *[U-<sup>13</sup>C]Glucose* |  |  |
| --- | --- | --- | --- |
|  | *Control* | *LPS* | *P-value* |
| **Glutamate** | 36.7±4.9 | 51.6±13 | 0.14, *n.s.* |
| **Glutamine** | **37.3±8.1** | **60.3±10** | **0.03, \*** |
| **Taurine** | **61.0±6.5** | **49.4±5.3** | **0.03, \*** |
| **GABA** | **1.3±0.3** | **3.6±1.8** | **0.01, \*** |

## References

Aldana, B. I. (2019). Microglia-Specific Metabolic Changes in Neurodegeneration. J Mol Biol, 431(9), 1830–1842. 10.1016/j.jmb.2019.03.006

Andersen, J. V., Christensen, S. K., Westi, E. W., Diaz-delCastillo, M., Tanila, H., Schousboe, A., Aldana, B. I., & Waagepetersen, H. S. (2021). Deficient astrocyte metabolism impairs glutamine synthesis and neurotransmitter homeostasis in a mouse model of Alzheimer’s disease. Neurobiol Dis, 148, 105198. 10.1016/j.nbd.2020.105198

Andersen, J. V., Jakobsen, E., Westi, E. W., Lie, M. E. K., Voss, C. M., Aldana, B. I., Schousboe, A., Wellendorph, P., Bak, L. K., Pinborg, L. H., & Waagepetersen, H. S. (2020). Extensive astrocyte metabolism of gamma-aminobutyric acid (GABA) sustains glutamine synthesis in the mammalian cerebral cortex. Glia, 68(12), 2601–2612. 10.1002/glia.23872

Andersen, J. V., Nissen, J. D., Christensen, S. K., Markussen, K. H., & Waagepetersen, H. S. (2017). Impaired Hippocampal Glutamate and Glutamine Metabolism in the db/db Mouse Model of Type 2 Diabetes Mellitus. Neural Plast, 2017, 2107084. 10.1155/2017/2107084

Andersen, J. V., & Schousboe, A. (2023). Glial Glutamine Homeostasis in Health and Disease. Neurochem Res, 48(4), 1100–1128. 10.1007/s11064-022-03771-1

Andersen, J. V., Schousboe, A., & Verkhratsky, A. (2022). Astrocyte energy and neurotransmitter metabolism in Alzheimer’s disease: Integration of the glutamate/GABA-glutamine cycle. Prog Neurobiol, 217, 102331. 10.1016/j.pneurobio.2022.102331

Andersen, J. V., Schousboe, A., & Wellendorph, P. (2023). Astrocytes regulate inhibitory neurotransmission through GABA uptake, metabolism, and recycling. Essays Biochem, 67(1), 77–91. 10.1042/ebc20220208

Andersen, J. V., Westi, E. W., Griem-Krey, N., Skotte, N. H., Schousboe, A., Aldana, B. I., & Wellendorph, P. (2024). Deletion of CaMKIIalpha disrupts glucose metabolism, glutamate uptake, and synaptic energetics in the cerebral cortex. J Neurochem, 168(5), 704–718. 10.1111/jnc.15814

Andersen, J. V., Westi, E. W., Jakobsen, E., Urruticoechea, N., Borges, K., & Aldana, B. I. (2021). Astrocyte metabolism of the medium-chain fatty acids octanoic acid and decanoic acid promotes GABA synthesis in neurons via elevated glutamine supply. Mol Brain, 14(1), 132. 10.1186/s13041-021-00842-2

Baik, S. H., Kang, S., Lee, W., Choi, H., Chung, S., Kim, J. I., & Mook-Jung, I. (2019). A Breakdown in Metabolic Reprogramming Causes Microglia Dysfunction in Alzheimer’s Disease. Cell Metab, 30(3), 493–507.e496. 10.1016/j.cmet.2019.06.005

Barger, S. W., Goodwin, M. E., Porter, M. M., & Beggs, M. L. (2007). Glutamate release from activated microglia requires the oxidative burst and lipid peroxidation. J Neurochem, 101(5), 1205–1213. 10.1111/j.1471-4159.2007.04487.x

Bernier, L. P., York, E. M., Kamyabi, A., Choi, H. B., Weilinger, N. L., & MacVicar, B. A. (2020). Microglial metabolic flexibility supports immune surveillance of the brain parenchyma. Nat Commun, 11(1), 1559. 10.1038/s41467-020-15267-z

Bernier, L. P., York, E. M., & MacVicar, B. A. (2020). Immunometabolism in the Brain: How Metabolism Shapes Microglial Function. Trends Neurosci, 43(11), 854–869. 10.1016/j.tins.2020.08.008

Bhandage, A. K., Jin, Z., Korol, S. V., Shen, Q., Pei, Y., Deng, Q., Espes, D., Carlsson, P. O., Kamali-Moghaddam, M., & Birnir, B. (2018). GABA Regulates Release of Inflammatory Cytokines From Peripheral Blood Mononuclear Cells and CD4(+) T Cells and Is Immunosuppressive in Type 1 Diabetes. EBioMedicine, 30, 283–294. 10.1016/j.ebiom.2018.03.019

Bhat, R., Axtell, R., Mitra, A., Miranda, M., Lock, C., Tsien, R. W., & Steinman, L. (2010). Inhibitory role for GABA in autoimmune inflammation. Proc Natl Acad Sci U S A, 107(6), 2580–2585. 10.1073/pnas.0915139107

Bi, D., Wen, L., Wu, Z., & Shen, Y. (2020). GABAergic dysfunction in excitatory and inhibitory (E/I) imbalance drives the pathogenesis of Alzheimer’s disease. Alzheimers Dement, 16(9), 1312–1329. 10.1002/alz.12088

Bradford, H. F., Ward, H. K., & Thomas, A. J. (1978). Glutamine--a major substrate for nerve endings. J Neurochem, 30(6), 1453–1459. 10.1111/j.1471-4159.1978.tb10477.x

Buck, M. D., O’Sullivan, D., & Pearce, E. L. (2015). T cell metabolism drives immunity. J Exp Med, 212(9), 1345–1360. 10.1084/jem.20151159

Butovsky, O., & Weiner, H. L. (2018). Microglial signatures and their role in health and disease. Nat Rev Neurosci, 19(10), 622–635. 10.1038/s41583-018-0057-5

Caputa, G., Castoldi, A., & Pearce, E. J. (2019). Metabolic adaptations of tissue-resident immune cells. Nat Immunol, 20(7), 793–801. 10.1038/s41590-019-0407-0

Fattorini, G., Catalano, M., Melone, M., Serpe, C., Bassi, S., Limatola, C., & Conti, F. (2020). Microglial expression of GAT-1 in the cerebral cortex. Glia, 68(3), 646–655. 10.1002/glia.23745

Frank, S., Burbach, G. J., Bonin, M., Walter, M., Streit, W., Bechmann, I., & Deller, T. (2008). TREM2 is upregulated in amyloid plaque-associated microglia in aged APP23 transgenic mice. Glia, 56(13), 1438–1447. 10.1002/glia.20710

Gimeno-Bayón, J., López-López, A., Rodríguez, M. J., & Mahy, N. (2014). Glucose pathways adaptation supports acquisition of activated microglia phenotype. J Neurosci Res, 92(6), 723–731. 10.1002/jnr.23356

Hanisch, U.-K., & Kettenmann, H. (2007). Microglia: active sensor and versatile effector cells in the normal and pathologic brain. Nature Neuroscience, 10(11), 1387–1394. 10.1038/nn1997

He, Y., Taylor, N., Yao, X., & Bhattacharya, A. (2021). Mouse primary microglia respond differently to LPS and poly(I:C) in vitro. Sci Rep, 11(1), 10447. 10.1038/s41598-021-89777-1

He, Y., Taylor, N., Yao, X., & Bhattacharya, A. (2021). Mouse primary microglia respond differently to LPS and poly(I:C) in vitro. Scientific Reports, 11(1), 10447. 10.1038/s41598-021-89777-1

Hines, D. J., Choi, H. B., Hines, R. M., Phillips, A. G., & MacVicar, B. A. (2013). Prevention of LPS-induced microglia activation, cytokine production and sickness behavior with TLR4 receptor interfering peptides. PLoS One, 8(3), e60388. 10.1371/journal.pone.0060388

Horvath, R. J., Nutile-McMenemy, N., Alkaitis, M. S., & Deleo, J. A. (2008). Differential migration, LPS-induced cytokine, chemokine, and NO expression in immortalized BV-2 and HAPI cell lines and primary microglial cultures. J Neurochem, 107(2), 557–569. 10.1111/j.1471-4159.2008.05633.x

Hu, Y., Mai, W., Chen, L., Cao, K., Zhang, B., Zhang, Z., Liu, Y., Lou, H., Duan, S., & Gao, Z. (2020). mTOR-mediated metabolic reprogramming shapes distinct microglia functions in response to lipopolysaccharide and ATP. Glia, 68(5), 1031–1045. 10.1002/glia.23760

Huang, Q., Wang, Y., Chen, S., & Liang, F. (2024). Glycometabolic Reprogramming of Microglia in Neurodegenerative Diseases: Insights from Neuroinflammation. Aging Dis, 15(3), 1155–1175. 10.14336/ad.2023.0807

Imai, Y., Ibata, I., Ito, D., Ohsawa, K., & Kohsaka, S. (1996). A novel gene iba1 in the major histocompatibility complex class III region encoding an EF hand protein expressed in a monocytic lineage. Biochem Biophys Res Commun, 224(3), 855–862. 10.1006/bbrc.1996.1112

Ito, D., Imai, Y., Ohsawa, K., Nakajima, K., Fukuuchi, Y., & Kohsaka, S. (1998). Microglia-specific localisation of a novel calcium binding protein, Iba1. Brain Res Mol Brain Res, 57(1), 1–9. 10.1016/s0169-328x(98)00040-0

Jin, L. W., Horiuchi, M., Wulff, H., Liu, X. B., Cortopassi, G. A., Erickson, J. D., & Maezawa, I. (2015). Dysregulation of glutamine transporter SNAT1 in Rett syndrome microglia: a mechanism for mitochondrial dysfunction and neurotoxicity. J Neurosci, 35(6), 2516–2529. 10.1523/JNEUROSCI.2778-14.2015

Kang, S., Liu, L., Wang, T., Cannon, M., Lin, P., Fan, T. W., Scott, D. A., Wu, H. J., Lane, A. N., & Wang, R. (2022). GAB functions as a bioenergetic and signalling gatekeeper to control T cell inflammation. Nat Metab, 4(10), 1322–1335. 10.1038/s42255-022-00638-1

Kettenmann, H., Hanisch, U. K., Noda, M., & Verkhratsky, A. (2011). Physiology of microglia. Physiol Rev, 91(2), 461–553. 10.1152/physrev.00011.2010

Kurki, S. N., Srinivasan, R., Laine, J., Virtanen, M. A., Ala-Kurikka, T., Voipio, J., & Kaila, K. (2023). Acute neuroinflammation leads to disruption of neuronal chloride regulation and consequent hyperexcitability in the dentate gyrus. Cell Rep, 42(11), 113379. 10.1016/j.celrep.2023.113379

Lauro, C., & Limatola, C. (2020). Metabolic Reprograming of Microglia in the Regulation of the Innate Inflammatory Response. Front Immunol, 11, 493. 10.3389/fimmu.2020.00493

Lian, H., Roy, E., & Zheng, H. (2016). Protocol for Primary Microglial Culture Preparation. Bio Protoc, 6(21). 10.21769/BioProtoc.1989

Liu, W., Taso, O., Wang, R., Bayram, S., Graham, A. C., Garcia-Reitboeck, P., Mallach, A., Andrews, W. D., Piers, T. M., Botia, J. A., Pocock, J. M., Cummings, D. M., Hardy, J., Edwards, F. A., & Salih, D. A. (2020). Trem2 promotes anti-inflammatory responses in microglia and is suppressed under pro-inflammatory conditions. Hum Mol Genet, 29(19), 3224–3248. 10.1093/hmg/ddaa209

Lu, J., Zhou, W., Dou, F., Wang, C., & Yu, Z. (2021). TRPV1 sustains microglial metabolic reprogramming in Alzheimer’s disease. EMBO Rep, 22(6), e52013. 10.15252/embr.202052013

Lund, S., Christensen, K. V., Hedtjarn, M., Mortensen, A. L., Hagberg, H., Falsig, J., Hasseldam, H., Schrattenholz, A., Porzgen, P., & Leist, M. (2006). The dynamics of the LPS triggered inflammatory response of murine microglia under different culture and in vivo conditions. J Neuroimmunol, 180(1-2), 71–87. 10.1016/j.jneuroim.2006.07.007

Miller, A. H., Haroon, E., Raison, C. L., & Felger, J. C. (2013). Cytokine targets in the brain: impact on neurotransmitters and neurocircuits. Depress Anxiety, 30(4), 297–306. 10.1002/da.22084

Mills, E. L., Kelly, B., Logan, A., Costa, A. S. H., Varma, M., Bryant, C. E., Tourlomousis, P., Dabritz, J. H. M., Gottlieb, E., Latorre, I., Corr, S. C., McManus, G., Ryan, D., Jacobs, H. T., Szibor, M., Xavier, R. J., Braun, T., Frezza, C., Murphy, M. P., & O’Neill, L. A. (2016). Succinate Dehydrogenase Supports Metabolic Repurposing of Mitochondria to Drive Inflammatory Macrophages. Cell, 167(2), 457–470 e413. 10.1016/j.cell.2016.08.064

Nair, S., Sobotka, K. S., Joshi, P., Gressens, P., Fleiss, B., Thornton, C., Mallard, C., & Hagberg, H. (2019). Lipopolysaccharide-induced alteration of mitochondrial morphology induces a metabolic shift in microglia modulating the inflammatory response in vitro and in vivo. Glia, 67(6), 1047–1061. 10.1002/glia.23587

Nayak, D., Roth, T. L., & McGavern, D. B. (2014). Microglia development and function. Annu Rev Immunol, 32, 367–402. 10.1146/annurev-immunol-032713-120240

Nimmerjahn, A., Kirchhoff, F., & Helmchen, F. (2005). Resting microglial cells are highly dynamic surveillants of brain parenchyma in vivo. Science, 308(5726), 1314–1318. 10.1126/science.1110647

O’Flaherty, L., Stapleton, P. P., Redmond, H. P., & Bouchier-Hayes, D. (1997). Dexamethasone and lipopolysaccharide regulation of taurine transport in Caco-2 cells. J Surg Res, 69(2), 331–336. 10.1006/jsre.1997.5067

Palsson-McDermott, E. M., & O’Neill, L. A. J. (2013). The Warburg effect then and now: From cancer to inflammatory diseases. BioEssays, 35(11), 965–973. 10.1002/bies.201300084

Park, E., Jia, J., Quinn, M. R., & Schuller-Levis, G. (2002). Taurine chloramine inhibits lymphocyte proliferation and decreases cytokine production in activated human leukocytes. Clin Immunol, 102(2), 179–184. 10.1006/clim.2001.5160

Persson, M., Brantefjord, M., Hansson, E., & Ronnback, L. (2005). Lipopolysaccharide increases microglial GLT-1 expression and glutamate uptake capacity in vitro by a mechanism dependent on TNF-alpha. Glia, 51(2), 111–120. 10.1002/glia.20191

Peruzzotti-Jametti, L., Willis, C. M., Hamel, R., Krzak, G., & Pluchino, S. (2021). Metabolic Control of Smoldering Neuroinflammation. Front Immunol, 12, 705920. 10.3389/fimmu.2021.705920

Qin, L., Li, G., Qian, X., Liu, Y., Wu, X., Liu, B., Hong, J. S., & Block, M. L. (2005). Interactive role of the toll-like receptor 4 and reactive oxygen species in LPS-induced microglia activation. Glia, 52(1), 78–84. 10.1002/glia.20225

Roqué, P. J., & Costa, L. G. (2017). Co-Culture of Neurons and Microglia. Curr Protoc Toxicol, 74, 11.24.11-11.24.17. 10.1002/cptx.32

Sahara, S., Yanagawa, Y., O’Leary, D. D., & Stevens, C. F. (2012). The fraction of cortical GABAergic neurons is constant from near the start of cortical neurogenesis to adulthood. J Neurosci, 32(14), 4755–4761. 10.1523/jneurosci.6412-11.2012

Salcedo, C., Wagner, A., Andersen, J. V., Vinten, K. T., Waagepetersen, H. S., Schousboe, A., Freude, K. K., & Aldana, B. I. (2021). Downregulation of GABA Transporter 3 (GAT3) is Associated with Deficient Oxidative GABA Metabolism in Human Induced Pluripotent Stem Cell-Derived Astrocytes in Alzheimer’s Disease. Neurochem Res, 46(10), 2676–2686. 10.1007/s11064-021-03276-3

Stratoulias, V., Venero, J. L., Tremblay, M. E., & Joseph, B. (2019). Microglial subtypes: diversity within the microglial community. Embo j, 38(17), e101997. 10.15252/embj.2019101997

Stuckey, D. J., Anthony, D. C., Lowe, J. P., Miller, J., Palm, W. M., Styles, P., Perry, V. H., Blamire, A. M., & Sibson, N. R. (2005). Detection of the inhibitory neurotransmitter GABA in macrophages by magnetic resonance spectroscopy. J Leukoc Biol, 78(2), 393–400. 10.1189/jlb.1203604

Takeda, H., Murakami, S., Liu, Z., Sawa, T., Takahashi, M., Izumi, Y., Bamba, T., Sato, H., Akaike, T., Sekine, H., & Motohashi, H. (2023). Sulfur metabolic response in macrophage limits excessive inflammatory response by creating a negative feedback loop. Redox Biol, 65, 102834. 10.1016/j.redox.2023.102834

Tannahill, G. M., Curtis, A. M., Adamik, J., Palsson-McDermott, E. M., McGettrick, A. F., Goel, G., Frezza, C., Bernard, N. J., Kelly, B., Foley, N. H., Zheng, L., Gardet, A., Tong, Z., Jany, S. S., Corr, S. C., Haneklaus, M., Caffrey, B. E., Pierce, K., Walmsley, S., … O’Neill, L. A. (2013). Succinate is an inflammatory signal that induces IL-1beta through HIF-1alpha. Nature, 496(7444), 238–242. 10.1038/nature11986

Tu, D., Gao, Y., Yang, R., Guan, T., Hong, J. S., & Gao, H. M. (2019). The pentose phosphate pathway regulates chronic neuroinflammation and dopaminergic neurodegeneration. J Neuroinflammation, 16(1), 255. 10.1186/s12974-019-1659-1

Ulland, T. K., Song, W. M., Huang, S. C., Ulrich, J. D., Sergushichev, A., Beatty, W. L., Loboda, A. A., Zhou, Y., Cairns, N. J., Kambal, A., Loginicheva, E., Gilfillan, S., Cella, M., Virgin, H. W., Unanue, E. R., Wang, Y., Artyomov, M. N., Holtzman, D. M., & Colonna, M. (2017). TREM2 Maintains Microglial Metabolic Fitness in Alzheimer’s Disease. Cell, 170(4), 649–663 e613. 10.1016/j.cell.2017.07.023

Van den Bossche, J., O’Neill, L. A., & Menon, D. (2017). Macrophage Immunometabolism: Where Are We (Going)? Trends Immunol, 38(6), 395–406. 10.1016/j.it.2017.03.001

Viola, A., Munari, F., Sanchez-Rodriguez, R., Scolaro, T., & Castegna, A. (2019). The Metabolic Signature of Macrophage Responses. Front Immunol, 10, 1462. 10.3389/fimmu.2019.01462

Voloboueva, L. A., Emery, J. F., Sun, X., & Giffard, R. G. (2013). Inflammatory response of microglial BV-2 cells includes a glycolytic shift and is modulated by mitochondrial glucose-regulated protein 75/mortalin. FEBS Lett, 587(6), 756–762. 10.1016/j.febslet.2013.01.067

Wake, H., Moorhouse, A. J., Jinno, S., Kohsaka, S., & Nabekura, J. (2009). Resting microglia directly monitor the functional state of synapses in vivo and determine the fate of ischemic terminals. J Neurosci, 29(13), 3974–3980. 10.1523/JNEUROSCI.4363-08.2009

Walls, A. B., Bak, L. K., Sonnewald, U., Schousboe, A., & Waagepetersen, H. S. (2014). Metabolic Mapping of Astrocytes and Neurons in Culture Using Stable Isotopes and Gas Chromatography-Mass Spectrometry (GC-MS). In J. Hirrlinger & H. S. Waagepetersen (Eds.), Brain Energy Metabolism (pp. 73–105). Springer New York. 10.1007/978-1-4939-1059-5_4

Walls, A. B., Bak, L. K., Sonnewald, U., Schousboe, A., & Waagepetersen, H. S. (2014). Metabolic Mapping of Astrocytes and Neurons in Culture Using Stable Isotopes and Gas Chromatography-Mass Spectrometry (GC-MS). In J. Hirrlinger & H. S. Waagepetersen (Eds.), Brain Energy Metabolism. Neuromethods, vol 90. Humana Press.

Walter, S., Letiembre, M., Liu, Y., Heine, H., Penke, B., Hao, W., Bode, B., Manietta, N., Walter, J., Schulz-Schuffer, W., & Fassbender, K. (2007). Role of the toll-like receptor 4 in neuroinflammation in Alzheimer’s disease. Cell Physiol Biochem, 20(6), 947–956. 10.1159/000110455

Westi, E. W., Jakobsen, E., Voss, C. M., Bak, L. K., Pinborg, L. H., Aldana, B. I., & Andersen, J. V. (2022). Divergent Cellular Energetics, Glutamate Metabolism, and Mitochondrial Function Between Human and Mouse Cerebral Cortex. Mol Neurobiol, 59(12), 7495–7512. 10.1007/s12035-022-03053-5

Yamada, D., Kawabe, K., Tosa, I., Tsukamoto, S., Nakazato, R., Kou, M., Fujikawa, K., Nakamura, S., Ono, M., Oohashi, T., Kaneko, M., Go, S., Hinoi, E., Yoneda, Y., & Takarada, T. (2019). Inhibition of the glutamine transporter SNAT1 confers neuroprotection in mice by modulating the mTOR-autophagy system. Commun Biol, 2, 346. 10.1038/s42003-019-0582-4

York, E. M., Zhang, J., Choi, H. B., & MacVicar, B. A. (2021). Neuroinflammatory inhibition of synaptic long-term potentiation requires immunometabolic reprogramming of microglia. Glia, 69(3), 567–578. 10.1002/glia.23913

Zhang, Z., Li, M., Li, X., Feng, Z., Luo, G., Wang, Y., & Gao, X. (2024). Glutamine metabolism modulates microglial NLRP3 inflammasome activity through mitophagy in Alzheimer’s disease. J Neuroinflammation, 21(1), 261. 10.1186/s12974-024-03254-w

Zhou, Y., & Danbolt, N. C. (2013). GABA and Glutamate Transporters in Brain. Front Endocrinol (Lausanne), 4, 165. 10.3389/fendo.2013.00165

